# Capturing trophectoderm-like stem cells enables step-wisely remodeling of placental development

**DOI:** 10.1101/2025.08.25.672082

**Authors:** Xinyi Jia, Bing Peng, Hongjin Zhao, Chunhui Wang, Wei Tao, Peng Du

## Abstract

The trophectoderm produced from totipotent blastomeres initiates trophoblast development, while placental deficiencies can cause pregnancy disorders. Yet, a culture system that fully recapitulates the entire placenta development is still lacking, greatly limiting related studies. Here, we captured mouse trophectoderm-like stem cells (TELSCs), which can give rise to all trophoblast lineages and be applied to generate trophoblast organoids. We achieved the induction and maintenance of TELSCs from totipotent blastomere-like stem cells or early embryos through a Hippo-YAP/Notch-to-TGFβ1 signaling switch. At the molecular level, TELSCs resemble E4.5 trophectoderm and are distinct from all previously known trophoblast-like stem cells. Functionally, TELSCs can generate all trophoblast lineages in both teratoma and chimera assays. We further applied TELSCs to generate trophoblast organoids containing various mature trophoblasts and a self-renewing extraembryonic ectoderm (ExE)-like progenitor population. Interestingly, we observed transiently formed rosette-like structures that rely on *Itgb1*, which are essential to induce ExE-like progenitors and to generate organoids eventually. Thus, the capture of TELSCs enables comprehensive insights into placental development.

**Highlights:**

- TELSCs were robustly induced and long-term maintained from TBLCs and early embryos using a newly developed Hippo-YAP/Notch-to-TGFβ1 signaling switch strategy
- TELSCs resemble E4.5 trophectoderm at the molecular level and generate all trophoblast lineages both *in vivo* and *in vitro* functionally
- TELSCs can be applied to generate trophoblast organoids containing multiple mature subtypes and a self-renewing ExE-like population
- Rosette structure formation that relies on *Itgb1* is critical to induce ExE-like progenitors and to eventually form trophoblast organoids

## Introduction

During murine embryonic development, trophoblasts originate from the outer cells of compacted embryos at the eight-cell stage ^1–6^. By embryonic day 3.5 (E3.5), the trophectoderm (TE) and inner cell mass (ICM) specification has been completed ^7–10^, culminating in embryo implantation into the uterus through the mural trophectoderm around E4.5 ^11,12^. The mural TE undergoes differentiation into trophoblast giant cells (TGCs), while the polar TE proliferates, giving rise to the extraembryonic ectoderm (ExE) and ectoplacental cone (EPC), consisting of TGCs and spongiotrophoblasts (SpTs) ^12–15^. Around E8.5, the chorionic epithelium derived from the ExE contacts the fetal mesoderm and differentiates into two layers of syncytiotrophoblasts (SynTs). Subsequently, the placental vascular network develops, forming the labyrinth, a densely packed structure ^16–18^. The mature placenta, around E12.5, comprises outer TGCs, SpTs, and inner labyrinth trophoblasts, serving as a transient essential organ facilitating efficient fetal-maternal communication for successful embryo development and pregnancy. Given the prevalence of developmental disorders that cause pregnancy failure in humans ^19^, comprehending the developmental program of trophoblast lineages and the placenta is crucial for reproductive and regenerative medicine.

*In vitro,* both human and mouse trophoblast stem cells (TSCs) have been derived from TE cells of blastocysts or ExE of post-implantation embryos ^20^, while under long-term culturing under defined conditions such as in TSC or TX medium, these mouse TSCs closely resemble trophoblast progenitors of E5.5-E7.5 embryos ^21^. Notably, recent studies have shown that naïve, or even primed human iPSCs or embryonic stem cells (ESCs) can be induced to generate TSCs ^22,23^, while in contrast, mouse pluripotent ESCs cannot be induced to form TSCs, suggesting that human pluripotent iPSCs/ESCs possess much higher plasticity than mouse ESCs. Particularly, quite recently, a novel type of so-called trophectoderm stem cells (TESCs) has been captured in a newly developed medium supplemented with Activin, IL11, BMP7, 8-Br cAMP and LPA, since transcriptomic comparison suggested that these cells may possess certain features of polar TE cells ^24^. However, a detailed comparison between TESCs and *in vivo* trophectoderm at different developmental stages is still required to clearly classify the status of TESCs.

However, the lack of an experimental model that accurately simulates *in vivo* conditions has limited our knowledge of placental development. Recently, human trophoblast organoids derived from placental villous tissue and trophoblast stem cells (TSCs), either directly isolated *in vivo* or differentiated from human embryonic stem cells (ESCs) *in vitro*, have been established as tools for *in vitro* studies of placental development and disease ^25–27^. However, these models exhibit an inside-out villous architecture and contain limited cell types, differing from native placental villi. Similarly, a recent study has demonstrated that mouse trophoblast organoids can also be derived from placentas or TSCs ^28^. However, two separate media are required to support organoid proliferation and differentiation, and the differentiation culture system lacks syncytiotrophoblast layer II (SynTII) cell types. Furthermore, both human and mouse TSC-to-trophoblast organoid formation systems only briefly recapitulate post-implantation placental development. The initiation of TE fate specification and the pre-implantation TE state transitions still cannot be recaptured *in vitro*. Thus, a trophoblast differentiation system capable of mimicking the entire stepwise placental development process initiated from totipotent stem cells remains unavailable.

In this study, we captured a novel type of trophectoderm-like stem cells (TELSCs), which can be applied to the stepwise remodeling of the entire placental development. Based on the Hippo-Yap/Notch-to-TGFβ1 signaling switch we developed the “two-step” system, which enabled the robust induction and stable propagation of TELSCs from both *in vitro* cultured totipotent blastomere-like cells (TBLCs) ^29^ and directly from *in vivo* 8-cell-stage mouse embryos. Molecularly, TELSCs closely resemble the TE cells of E4.5 blastocysts at both transcriptomic and epigenetic levels, and are clearly distinct from conventional TSCs and recently reported TESCs. Remarkably, in mouse teratoma and chimera assays, we demonstrated that TELSCs were able to successfully produce all the placental trophoblast lineages at the single-cell level. Furthermore, TELSCs can be applied to readily generate trophoblast organoids with all mature trophoblasts and long-term passaging ability. Additionally, we identified a novel population of E5.5-6.5 ExE-like progenitor cells with a high cell proliferation rate, which enabled the coupled self-renewal and differentiation abilities of TELSC-derived organoids. Interestingly, during organoid formation, we observed a dynamic and transient morphological formation of rosette-like structures, relying on the key β1 signaling factor *Itgb1,* which was essential to induce ExE-like progenitors and eventually to generate organoids from TELSCs. This achievement not only deepens our understanding of stepwise trophoblast differentiation from totipotent stem cells, but also provides a robust *in vitro* system for comprehensively investigating crucial events governing placental development.

## Results

### Mouse TELSCs were induced and stably maintained from TBLCs and 8-cell blastomeres using a “two-step” culture system

The placenta, comprising diverse trophoblast lineages derived from trophectoderm, plays a pivotal role in mediating fetal-maternal communication during pregnancy, and placental deficiency is implicated in various human fertility disorders. However, the lack of *in vitro* trophoblast culture and differentiation systems, particularly for the pre-implantation stage, has greatly impeded our understanding of placental development and related diseases to date. Recently, a novel kind of mouse totipotent blastomere-like cells (TBLCs) closely resembling 2-/4-cell blastomeres, which can produce various embryonic and extraembryonic lineages including mature trophoblasts, has been captured and stably maintained *in vitro*^29,30^. Consequently, TBLCs serve as ideal “seed cells” for establishing a trophoblast differentiation system originating from totipotent stem cells.

We first plated ESCs and TBLCs into classical serum-containing TS medium ^20^ supplemented with various factors known to induce trophoblast lineages, such as FGF4 ^21^, Activin A ^31^, TGFβ1 ^32^, and BMP4 ^33^. After three days, TBLCs exhibited significant morphological changes, forming tight epithelium-like colonies characteristic of trophoblast cells specifically in the FGF4-containing TS medium, but not in other media. Notably, ESCs did not respond similarly (**Figures 1A, 1B and S1A**). Fluorescence-activated cell sorting (FACS) analysis on the typical TE-specific markers, including CDX2 and CD40, revealed a notable population of CDX2+/CD40+ TE-like cells, constituting approximately up to 14% of TBLC-derived cells. While pluripotent ESCs cultured under the same conditions failed to generate a similar cell population (**Figure 1C, S1B and S1C**). Further immunostaining analysis confirmed the successful induction of TE-like cells from TBLCs at the protein level (**Figure 1D**). Thus, TE-like cells can be easily and efficiently induced from TBLCs in TS medium, but not from pluripotent ESCs, and we name these transiently induced TE-like cells as trophectoderm-like cells (TELCs).

**Figure 1.**
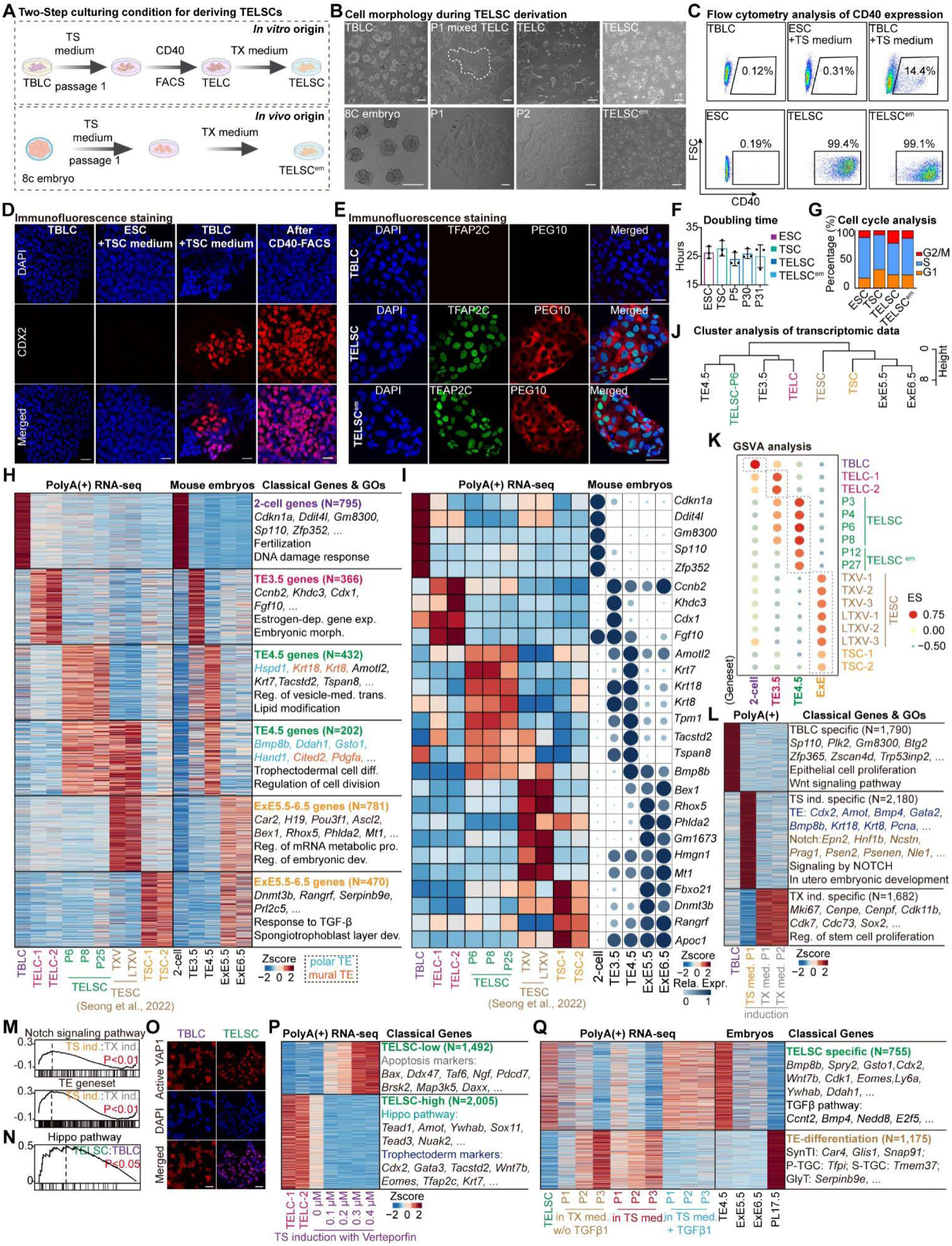
Capturing trophectoderm-like stem cells (TELSCs) with pre-implantation E4.5 TE features using a “two-step” culture system. (A) The diagram illustrating the process to obtain TELSCs (upper panel) and TELSC^em^s (lower panel). (B) The morphology of TELSCs (upper panel) and TELSC^em^s at different stage (lower panel). Scale bars, 100 µm. (C) FACS analysis of the percentage of CD40^+^ cells from TBLCs, as well as ESCs and TBLCs cultured in TS medium, using V6.5 cell line. (D) Immunofluorescence staining of CDX2 in CD40^+^ cells from ESCs and ESCs cultured in TS medium, as well as TBLCs cultured in TS medium before and after CD40 FACS purification. (E) Immunofluorescence staining of TFAP2C and PEG10 in TBLCs, TELSCs and TELSC^em^s. Scale bars, 50 μm. (F) Cell doubling time of ESCs, TSCs, TELSCs and TELSC^em^s at different passages. (G) Cell cycle analysis of ESCs, TSCs, TELSCs and TELSC^em^s. (H) Heatmap indicating the relative expression of differentially expressed genes related with mouse embryos^43–45^ in TBLCs, TELCs, TELSCs, TSCs and trophectoderm stem cells (TESCs). Transcriptomic data of TESCs is from ^46^. The enrichment of Gene Ontology (GO) terms of these genes is shown on the right. Dep., dependent; exp., expression; reg., regulation; trans., transport; diff.,differentiation; pro., process; dev., development. (I) Heatmap indicating the relative expression of characteristic genes in TBLCs, TELCs, TELSCs, TESCs and TSCs. Bubble chart showing the relative expression of these genes in mouse embryos. (J) Hierarchical clustering analysis of TELCs, TELSCs, TSCs, TESCs and mouse embryos data. (K) Gene Set Variation Analysis (GSVA) analysis of TBLCs, TELCs, TELSCs, TELSC^em^s, TESCs, TSCs and mouse embryos data. (L) Heatmap indicating the relative expression of differentially expressed genes of TBLCs and TBLCs induction in TS medium and TX medium. (M) Gene Set Enrichment Analysis (GSEA) analysis of TBLCs induction in TS medium and TX medium based on Notch signaling pathway and TE geneset. (N) GSEA of TELSCs versus TBLC ssamples based on Hippo pathway geneset. (O) Immunofluorescence staining of active-YAP1 in TELSCs and TBLCs. Scale bars, 50 µm. (P) Heatmap indicating the differentially expressed genes of TELCs and TBLCs induction in TS medium plus verteporfin. The representative genes and enrichment of GO terms of these genes is shown. (Q) Heatmap indicating the differentially expressed genes of TELSCs, TBLCs induction in TX medium withdraw TGFβ1, in TS medium and in TS medium plus TGFβ1. Heatmap on the right demonstrating the expression of each cluster in mouse embryos. The representative genes and enrichment of GO terms of these genes is shown.

However, these TELCs could not sustain a homogeneous morphology or undergo long-term passaging in the serum-containing TS medium (**Figure S1F**). Instead, they were stably maintained with stable TE-like morphology for more than 30 passages in the recently reported serum-free TX medium ^21^ (**Figures 1A and 1B**). Utilizing CD40-based FACS analysis, we observed that more than 99% of cells remained positive across various passages (**Figure 1C**), suggesting their steady and homogeneous cell status, which can be further confirmed by Western blot and immunostaining analysis by employing antibodies against TE-specific markers, including CDX2, EOMES, TFAP2C, and PEG10 (**Figure 1E, 1J and S1G**). We named these cells stably maintained in TX medium as trophectoderm-like stem cells (TELSCs).

Similarly, we plated embryos at different developmental stages in the same “two-step” culturing system and found that 8-cell embryos could give rise to homogeneous cells with TE morphology, which could be stably maintained over long-term passages (referred to as TELSC^em^; **Figures 1A, 1B and S1H**). FACS analysis revealed that over 99% of these embryo-derived cells were CD40-positive (**Figures 1C and S1I**), and immunostaining confirmed robust expression of key TE lineage markers, including CDX2, EOMES, HAND1, TFAP2C, and PEG10 (**Figures 1E and S1J**). Notably, TELSCs derived from both TBLCs and 8-cell embryos display rapid self-renewal ability, with a much higher proliferation rate than conventional TSCs, even after 30 passages (**Figure 1F**). While cell cycle analysis revealed that TELSCs maintained a typical active stem cell-like cycle distribution, characterized by an increased G2/M phase and a reduced S phase compared to pluripotent ESCs (**Figures 1G and S1K**).

Above all, we developed a “two-step” TS-TX culture strategy that enables the efficient and reproducible derivation of a novel type of TELSCs with typical TE features and a fast self-renewal ability from mouse TBLCs or 8-cell embryos.

### TELSCs are distinct from known TESCs/TSCs and are close to pre-implantation E4.5 TE cells at the transcriptomic level

Subsequently, we aimed to comprehensively characterize the molecular attributes of TELSCs stably maintained in TX medium, as well as transiently-induced TELCs in TS medium. RNA-seq was performed on these cells, and transcriptomic comparison clearly showed that compared to the original TBLCs, a total of 795 totipotent genes, including *Gm8300*, *Zfp352*, *Ddit4l*, and *Sp100*, were uniformly silenced in TELCs and TELSCs. While in TELCs, 366 genes represented by *Ccnb2*, *Cdx1*, and *Fgf10*, which are particularly enriched in TE cells of E3.5 embryos (TE3.5) and are related to “Estrogen-dependent gene expression”, were dynamically and specially activated. Furthermore, a large group of genes, such as *Amotf2*, *Krt7/8/18*, *Tspan8, Pdgfa, Cited2, Hand1* and *Tacstd2*, which are preferentially enriched in TE cells of E4.5 embryos (TE4.5) and linked to “Lipid modification” and “trophectoderm cell differentiation” GO terms, exhibited specific induction and stable expression in TELSCs across various passages (**Figures 1H, 1I, and S1L-S1O; Table S1**). Gene set enrichment analysis (GSEA) further revealed that both TELCs and TELSCs exhibited significant enrichment of placental development-related pathways, underscoring their functional resemblance to *in vivo* TE lineages (**Figure S1P**).

A recent study claimed to capture a novel type of trophectoderm stem cells (TESCs), which resembled E4.5 polar TE lineages ^24^. We next compared the TELCs and TELSCs we captured in this study, with TESCs, as well as the well-known TSCs cultured in both traditional serum-containing TS medium and optimized serum-free TX medium, at the transcriptomic level. We found that distinct from TELCs or TELSCs we captured in this study, TSCs exhibited obvious post-implantation E5.5-6.5 extraembryonic ectoderm (ExE) characteristics, with high expression of ExE-specific genes, such as *Prl2c5*, *Serpinb9e*, *Rangrf* and *Dnmt3b* (**Figures 1H and 1I**). Interestingly, although we indeed detected 202 genes, including *Cited2*, *Pdgfa*, *Hand1*, *Bmp8b* and *Gsto1*, were enriched in both TESCs and TELSCs cultured *in vitro*, as well as in E4.5 TE cells *in vivo*, TELSCs were particularly enriched with 781 genes, such as *Rho5*, *Phlda2*, *Hmgn1* and *Tcf7l2,* specifically expressed in E5.5-6.5 ExE tissues, which were not expressed in TELSCs. In contrast, there were 432 genes, including *Hspd1*, *Amotl2*, *Krt7/8/18* and *Tspan8*, specially enriched in E4.5 TE and TELSCs, which were not detected in TESCs (**Figures 1H and 1I**). Thus, we consider that TESCs are dual-featured cells possessing both pre-implantation E4.5 TE and post-implantation E5.5-6.5 ExE characteristics. While comparably, TELSCs represent a novel type of TE-like stem cells with pure E4.5 TE features.

To provide a precise comparative analysis of these distinct TE-like cell populations in relation to *in vivo* embryonic development, we performed clustering analysis based on whole-transcriptome profiles. This analysis clearly demonstrated that TELCs clustered closely with E3.5 TE cells, while TELSCs exhibited transcriptional similarities to E4.5 TE cells, reflecting their resemblance to pre-implantation TE lineages (**Figure 1J**). In contrast, both TESCs and TSCs showed global transcriptomic profiles more closely aligned with post-implantation ExE cells at E5.5–6.5 stages (**Figure 1J**). The above result was consistent with and further supported by gene set variation analysis (GSVA) based on differentially expressed genes (DEGs) in TE3.5-ExE6.5 cells (**Figure 1K**; **Table S2**). In summary, our findings indicate that distinct from previously reported TESCs or conventional TSCs, TELCs and TELSCs exhibit a resemblance to TE cells from E3.5 and E4.5 embryos during the pre-implantation stage, respectively, at the transcriptomic level.

### The Hippo-Yap /Notch signaling activation drives TELC specification from TBLCs or totipotent blastomeres in TS, but not TX, medium

Given that neither TX nor TS medium alone can be applied to induce and stably maintain TELSCs directly from TBLCs or early embryos (**Figure S1F and S1H**), we proposed that a cell signaling alteration underlying the “two-step” TS-to-TX medium switch is essential for TELSC induction and maintenance. To investigate this, we first performed RNA-seq analysis on TBLC-derived cells plated into TS or TX media alone for a few days. We found that these cells cannot retain the totipotent state with a dramatic decrease of totipotent-specific genes, such as *ZFP365*, *Sp100*, *Zscan4d*, *Btg2* and *Plk2*, compared to the original TBLCs. Instead, two large distinct groups of genes were dramatically induced in these cells. Particularly, TE-specific genes, including *Cdx2*, *Gata2*, and *Krt8/18,* were robustly activated in the TS medium, but not in the TX one, suggesting that certain cell signaling pathway(s) were induced to drive TE lineage specification specifically in TS medium (**Figure 1L**). While both GO enrichment analysis and GSEA showed a prominent activation of the Notch and Hippo-YAP signaling pathways, both of which are known to initiate the first TE fate specification^34^, represented by key markers, including *Epn2*, *Prag1*, and *Nle1* for Notch signaling, and *Tead4*, *Wwc*, and *Yap1* for Hippo signaling, under TS, but not TX, conditions (**Figures 1L-1N and S1Q-S1R**). Using antibodies against active-YAP1, we performed immunostaining analysis, and we detected active-YAP1 expression clearly localized in the nucleus of TELSCs, whereas it was weakly expressed in TBLCs (**Figure 1O**). We thus proposed that Hippo-YAP/Notch signaling pathways drive TELC induction from TBLCs or 8-cell embryos, which occurs specifically in TS, but not TX, medium.

To functionally verify this, we supplemented Verteporfin, a Hippo-YAP signaling inhibitor, into TS medium when we began to induce TELC induction from TBLCs. After 24 hours, the Verteporfin treatment obviously blocked the formation of TE-like colonies (**Figure S1S**). Subsequent RNA-seq analysis revealed that, compared to control samples, Verteporfin-treated cells failed to activate key genes associated with the Hippo pathway, such as *Tead1* and *Amot*, which eventually led to the failure of TELC specification and activation of typical TE-specific genes, including *Emoes*, *Cdx2* and *Gata3.* While Verteporfin treatment led to the upregulation of several apoptosis-related genes (**Figures 1P, S1T and S1U**), explaining the cell death phenotype. Thus, Hippo-YAP/Notch signaling are required for TELC induction in TS medium, closely recapitulating the *in vivo* mechanism of TE specification, which is yet absent in TX medium.

### The TGFβ1 signaling and ITS-X enabled the maintenance and rapid self-renewal of TELSCs, respectively, in TX but not TS medium

Since TELSCs cannot be stably maintained in TS medium alone, although it enabled TELC specification, we proposed that some key signaling pathway(s) present in TX medium, but absent in TS medium, are responsible for the maintenance of TELSCs. To test this, we first transferred TELSCs that had been long-term cultured in TX medium back into TS medium. We observed that these cells gradually lost viability and could not be maintained over time (**Figure S1V**). While we noted that TX medium contained two additional components: ITS-X supplement and TGFβ1, which were absent in TS medium and likely supported the maintenance of TELSCs. We then removed both or each component from the TX medium and found that all of which led to the failure of TELSC maintenance after three passages. Bulk RNA-seq analysis revealed that cells cultured in TX medium without ITS-X retained high expression of TE-specific markers, including *Cdx2*, *Gata2*, *Krt8/18,* enriched in TELSCs and a TE-like transcriptional profile, while instead the expression of another group of cell proliferation genes was clearly decreased (**Figures S1W and S1X**). Thus, ITS-X, as known to support cell proliferation, enabled the rapid proliferation and long-term self-renewal of TELSCs in TX medium, which did not impact TELSC identity.

In contrast, the removal of TGFβ1 caused the rapid decrease of typical TE-specific genes, such as *Cdx2*, *Bmp4/8*, *Klf5* and Ly6a, which were highly expressed in TELSCs. Instead, mature trophoblast lineage-specific genes, including *Car4, Glis1, Snap91* for SynTI, *Tfpi* for P-TGC, *Tmem37* for S-TGC, and *Serpinb9e* for GlyT, were dramatically induced, showing the differentiation fate of original TELSCs. Thus, TGFβ1 signaling, which was absent in TS medium, was essential for the maintenance of the TELSC state in TX medium (**Figure 1Q**). Above all, we proposed that TGFβ1 together with ITS-X eventually allowed the maintenance and rapid self-renewal of TELSCs for long-term passages.

To further validate this, we transferred TELSCs, stably maintained in TX medium, into TS medium supplemented with either TGFβ1 or ITS-X. After several passages, we noticed that the addition of TGFβ1, but not ITS-X, enabled TELSCs to be maintained with typical TE-like morphology for long-term passages (beyond passage 11) (**Figure S1V**). RNA-seq analysis showed that the addition of TGFβ1 was indeed able to retain the widespread expression of typical TE-associated genes, including *Cdx2, Eomes, Ly6a, and Bmp8b*, allowing the maintenance of TELSC identity in the TX medium. Comparably, the addition of ITS-X could not prevent the differentiation fate of TELSCs with widespread induction of mature trophoblast lineage-specific genes, such as *Car4* and*Tmem37* (**Figure 1Q**).

Collectively, we demonstrate that Hippo-YAP/Notch signaling activation enables the first TE fate decision from totipotent cells, including TBLCs and 8-cell blastomeres, to generate TELCs in the TS medium. After that, in the follow-up TX medium, TGFβ1 signaling enhances the TELC-to-TELSC transition and eventually enables TELSC maintenance, while ITS-X facilitates the stable and rapid proliferation of TELSCs for long-term passages.

### The unique epigenomic features of TELCs and TELSCs distinct from TSCs

We next thought to characterize the epigenetic features of TELCs and TELSCs, and we performed transposase-accessible chromatin sequencing (ATAC-seq), cleavage under targets and tagmentation (CUT&Tag) for histone modifications, and whole-genome bisulfite sequencing (WGBS) to assess chromatin accessibility, histone modification patterns, and DNA methylation status, respectively. Notably, ATAC-seq analysis targeting transcription start sites (TSSs) revealed ATAC-seq signals in the promoter regions of 239 totipotent genes, including *Zdbf2*, *Cd80,* and *Klf3*, which were highly expressed in TBLCs but exhibited a pronounced decrease in TELCs and complete silencing in TELSCs, indicating a distinct open status in TBLCs that evidently transitioned to a closed status in TELSCs (**Figures 2A, 2B and S2A**). Furthermore, in the promoters of 439 genes, such as *Amotl2*, *Cdx2* and *Wnt9a*, which are selectively activated in TELCs, a TELC-specific chromatin open status was evident. Lastly, in the promoters of 150 genes, including *Elf5*, *Hand1,* and *Mbp*, particularly expressed in TELSCs, ATAC-seq signals gradually opened during the transition from TBLCs to TELSCs (**Figures 2A and 2B**).

**Figure 2.**
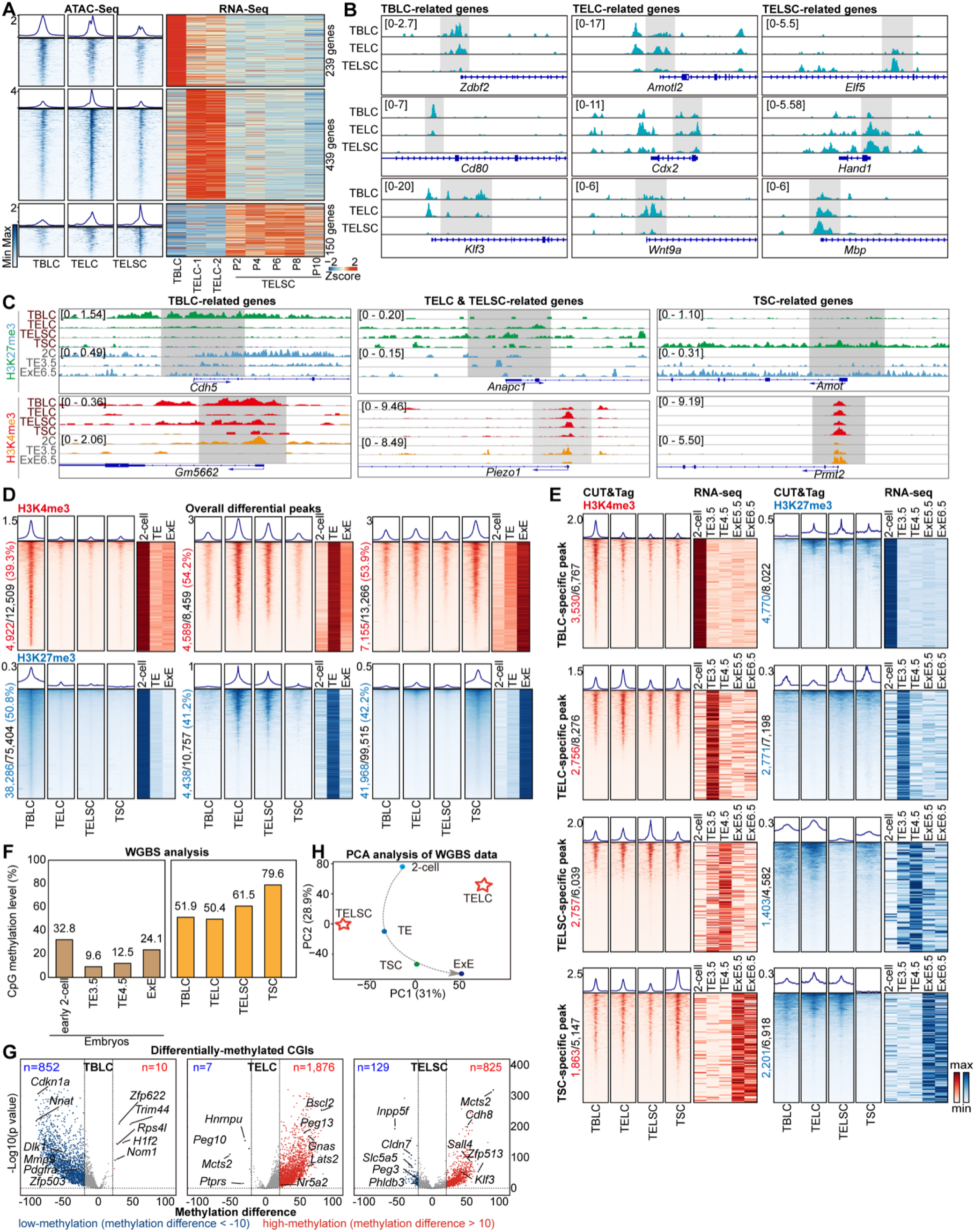
The unique epigenomic features of TELSCs. (A) ATAC-seq of TBLCs, TELCs and TELSCs. The left heatmaps show the open or closed peaks around transcription start sites (TSSs) in TBLCs, TELCs and TELSCs and the relative expression of related genes based on RNA-seq on the right heatmap (RNA-seq data of TBLCs is from ^29^. (B) IGV browser view displaying ATAC-seq signals of specific genes in TBLCs, TELCs and TELSCs. (C) IGV tracks showing H3K4me3 and H3K27me3 enrichment at representative marker genes across TBLCs, TELCs, TELSCs, and TSCs. (D) Left heatmap displaying the overall differential peaks across the genome of between TBLCs, TELCs, TELSCs and TSCs. Right heatmap showing the changes of differential peaks in mouse embryos ^35,36^. Black number, cell type specific peaks. Red or blue number, peaks comparable to mouse embryos data in vivo. Percentage, ratio of matching peaks (red or blue) to total (black). (E) Heatmaps displaying the overall H3K4me3 and H3K27me3 differential peaks across the genome of between TBLCs, TELCs, TELSCs and TSCs. Heatmaps on the right showing the changes in the expression of these genes in mouse embryos data *in vivo* ^43–45^. (F) Histograms showing the global average methylation levels of TBLCs, TELCs, and TELSCs and mouse embryos based on WGBS data. The global average methylation levels of mouse embryos is from ^43,44^. (G) WGBS analysis of TBLCs, TELCs and TELSCs. Volcano plots displaying the differentially methylated CpG islands in TBLCs, TELCs and TELSCs, respectively. For volcano plots, blue and red dots indicate low-methylation (methylation difference < −10) and high-methylation (methylation difference > 10) CpG islands of related genes with a p value < 0.05. WGBS data of TBLCs is from ^29^. (H) PCA analysis of TELCs, TELSCs, TSCs and mouse 2-cell, TE and ExE based on WGBS data. WGBS data of TSCs is from ^47^ and WGBS data of mouse TE and ExE is from ^43^.

Next, a genome-wide comparison of histone modification, including H3K4me3 and H3K27me3, across gene bodies with *in vivo* 2C, TE, and ExE6.5 stages ^35,36^ revealed that approximately half of the peaks in TBLCs, TELCs/TELSCs, and TSCs mirrored those in their corresponding *in vivo* counterparts (**Figures 2C and 2D**). In total, we identified 3,530 and 4,770 unique H3K4me3 and H3K27me3 peaks, respectively, in TBLCs, which were absent in trophoblast cells and correlated with genes highly expressed in TBLCs. TELCs and TELSCs harbored 2,756 and 2,757 H3K4me3 peaks, along with 2,771 and 1,403 H3K27me3 peaks, respectively, corresponding to genes upregulated in these populations. In TSCs, 1,863 H3K4me3-enriched regions and 2,201 regions with H3K27me3 depletion were identified, associated with genes predominantly expressed in ExE5.5/6.5 cells *in vivo* (**Figure 2E**). Representative totipotency markers such as *Gm5662* exhibited H3K4me3 enrichment specifically in TBLCs, while TELCs and TELSCs showed strong H3K4me3 signals at *Sirt1*, a gene essential for trophoblast stem cell differentiation and placental development ^37^. In contrast, *Amot*, a Hippo pathway member, was marked by H3K27me3 in TSCs but not in TELCs or TELSCs, suggesting a failure of TSCs to retain pre-implantation epigenetic features ^24^ (**Figure 2C**).

Finally, WGBS analysis clearly showed that TELCs and TELSCs do not exhibit increased global DNA methylation levels compared to TBLCs. In contrast, TSCs exhibit significantly higher global DNA methylation levels than TBLCs, TELCs, or TELSCs (**Figure 2F**). Additionally, the DNA methylation levels of many imprinted genes remain relatively stable without significant changes in TBLCs, TELCs, and TELSCs, which were much lower than those in TSCs (**Figure S2D**). Subsequent analysis of methylation status on CpG islands (CGIs) showed that CGIs on the promoters of 852 genes lacking DNA methylation in TBLCs displayed clear methylation in TELCs and TELSCs, suggesting a cell fate specification process from totipotent stem cells toward the trophectoderm lineage (**Figure 2G**). Additionally, we detected that TE-specific CGIs related to 1,876 genes (including *Peg13*, *Bscl2,* and *Gnas*) and 825 genes (containing *Cdh8*, *Sall4,* and *Zfp513*) were particularly methylated in TELCs and TELSCs, respectively. Additionally, we detected CGIs on the promoters of 7 genes (including *Peg10*, *Mcts2,* and *Hnrnpu*) and 129 genes (containing *Peg3*, *Inpp5f,* and *Phldb3*) that were demethylated and transcriptionally induced, particularly in TELCs and TELSCs, respectively (**Figure 2G**), illustrating TE-specific gene activation during TELC induction from TBLCs. PCA analysis using DNA methylation data revealed epigenetic similarities between TSCs and ExE6.5, while TELCs and TELSCs more closely resembled earlier stage TE **(Figure 2H)**.

In conclusion, the above comprehensive analysis elucidates the unique epigenetic features of TELCs and TELSCs that markedly differ from those of known TSCs.

### TELSCs specifically contribute to placental tissue with robust developmental potency to produce all trophoblast lineages

To precisely assess the developmental potency of TELSCs, we first performed *in vivo* embryo chimerism assays by injecting mCherry- or EGFP-labeled TELSCs or TSCs into donor 8-cell embryos. TELSCs consistently contributed to the TE lineage from early to fully hatched blastocyst stages, similar to TSCs (**Figures 3A and S3A**). Immunostaining analysis further confirmed that TELSCs efficiently and specifically integrated into the TE lineage, marked by CDX2 and KRT18, but not the embryonic epiblast lineage labeled by SOX2 (**Figures 3B, 3C and S3B-S3C**). At the later developmental stage around E13.5, we detected that TELSCs can widely contribute to placental tissues including both the junctional zone (JZ) and labyrinth (Lab) regions, but not yolk sac or fetal tissues (**Figure S3D**), showing the developmental specificity of TELSCs towards trophoblast lineages. Further immunohistochemistry analysis clearly demonstrated TELSCs produced various trophoblast lineages marked by the general trophoblast marker KRT7, as well as lineage-specific markers HAND1 and TPBPA, indicative of their differentiation into trophoblast giant cells (TGCs) and spongiotrophoblasts (SpTs), respectively (**Figure 3D**).

**Figure 3.**
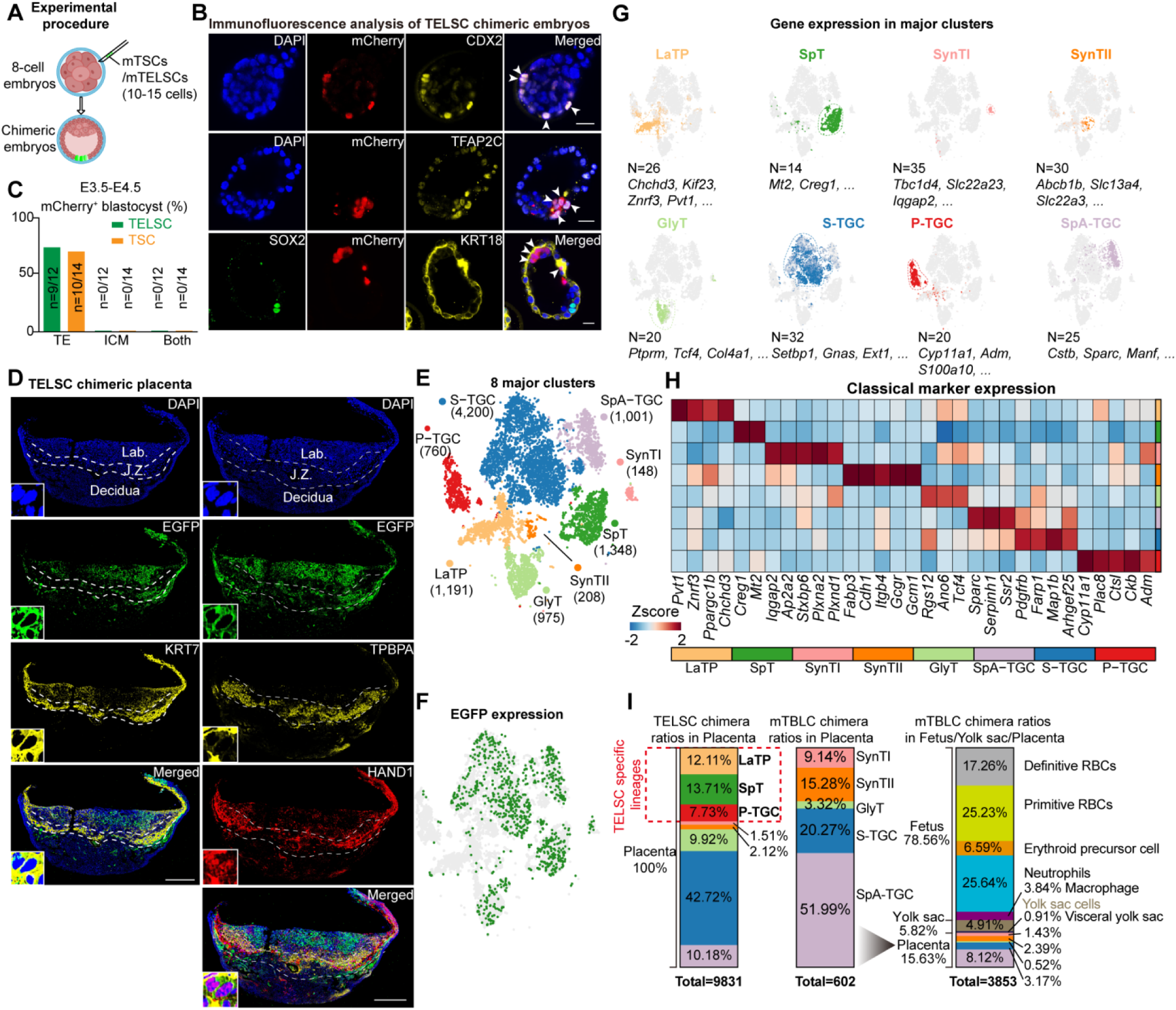
TELSCs exhibit robust *in vivo* developmental potential and full trophoblast lineage contribution. (A) Schematic of the experimental procedure to generate chimeric blastocysts. 10-15 EGFP-labeled TELSCs were injected into 8C-stage embryos. (B) Representative immunostaining of chimeric blastocysts injected with mCherry-labeled TELSCs. CDX2, TFAP2C and KRT18: TE-specific markers; SOX2: ICM-specific marker; white arrows, TELSCs contribute to TE. Scale bar, 20 µm. (C) Bar graph showing the chimerism ratio between TELSCs and TSCs. (D) Immunofluorescence analysis of placenta sections from E12.5 chimeric placentae. The placenta was stained with the KRT7, TPBPA, HAND1 and also EGFP antibody to amplify the fluorescent signal. Lab., labyrinth; J.Z., junctional zone. The insets show enlarged images of the white boxes. Scale bar, 1 mm. (E) t-SNE plot showing the 8 main trophoblastic clusters. (F) t-SNE plot showing EGFP expression in single EGFP^+^ cells from a chimeric placenta based on scRNA-seq. EGFP^+^ cells were sorted by flow cytometry. Green dots represent cells in which EGFP expression was detected in scRNA-seq. (G) t-SNE visualization showing marker gene expression in major clusters. (H) Heatmap showing the average expression of specific marker genes for each trophoblastic cluster. (I) Histogram indicating the proportion of cell types in TELSCs chimeric placentae and mTBLCs chimeric embryos.

To further precisely elucidate the differentiation potential of TELSCs, we performed scRNA-seq on placental tissues from E13.5 chimeric mice. The results provide clear and direct evidence that TELSCs contribute to all the eight trophoblast lineages reported to date, including labyrinth trophoblast progenitors (LaTPs), trophoblast glycogen cells (GlyTs), sinusoidal trophoblast giant cells (S-TGCs), spiral artery-associated trophoblast giant cells (SpA-TGCs), parietal trophoblast giant cells (P-TGCs), spongiotrophoblast cells (SpTs), and syncytiotrophoblasts layer I (SynTIs) and layer II (SynTIIs) (**Figures 3E-3H and S3E**). Notably, TELSCs showed substantial contribution to SpA-TGCs, a specialized trophoblast subtype essential for forming the junctional zone and mediating maternal–fetal communication (**Figures 3E and 3G**). While we further compared the cell lineage contribution of original TBLCs with that of TELSCs in the same chimeric assay^30^, we found that compared to TBLCs, which can widely contribute to various embryonic and extraembryonic cells, TELSCs specifically generate trophoblast lineages, but not any other embryonic cell types, therefore proving the very specific developmental potency of TELSCs, which eventually led to the widespread contribution of TELSCs to all trophoblast lineages (**Figure 3I**).

Nevertheless, for the very first time, our study directly and comprehensively demonstrates that TELSCs are able to produce all the trophoblast lineages *in vivo* at the single-cell resolution, since previous studies on TSCs, mainly based on immunostaining analysis, could not provide such definitive evidence, which therefore provides new criteria to precisely evaluate and compare the cell lineage contributions of various kinds of trophoblast-like stem cells.

### TELSCs exhibit superior trophoblast differentiation capacity compared to TSCs in the mouse teratoma assay

Beyond the above chimerism assay, we also conducted a teratoma analysis by injecting the same amount of TELSCs, TSCs, and mouse embryonic fibroblasts (MEFs, as a negative control) into immunodeficient nude mice, respectively (**Figure 4A**). We observed that both TSCs and TELSCs can efficiently form teratoma tissues, which can cause hemorrhagic lesions characterized by large blood-filled lacunae, but MEFs cannot (**Figures 4B and S4A**). Further histological analysis of these lesions revealed a typical trophoblastic hemorrhagic structure with ELF5-positive trophoblast giant cells (TGCs) that were differentiated from TELSCs or TSCs (**Figure 4C**), clearly showing the invasive properties of TGCs. An immunohistofluorescence assay further confirmed the presence of trophoblasts expressing KRT7 and PEG10, including SpTs and TGCs marked by TPBPA and PRL, respectively, in both TELSC- and TSC-derived teratomas (**Figure S4B**), demonstrating the trophoblast developmental potency of both TELSCs and TSCs.

**Figure 4.**
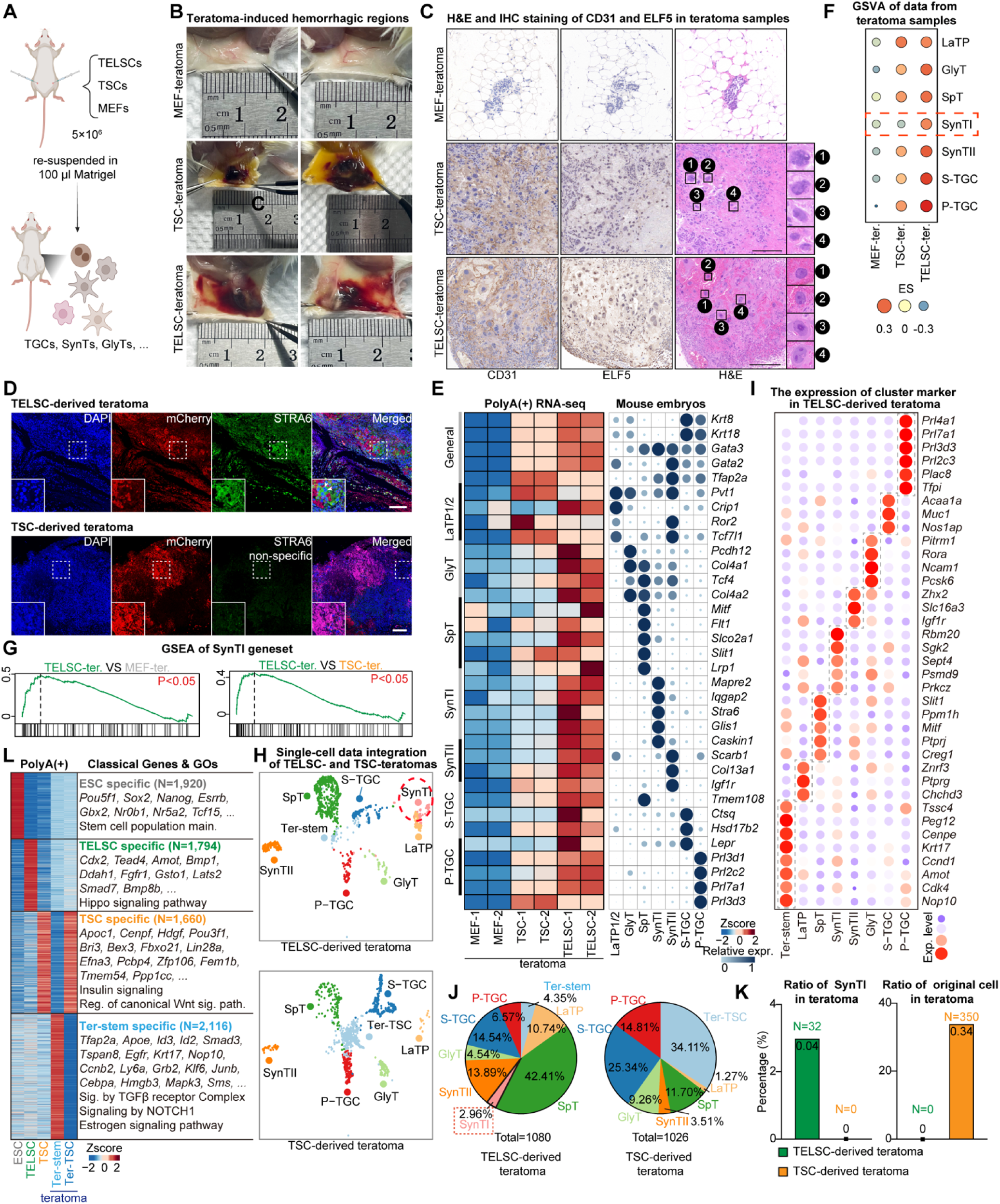
TELSCs exhibit enhanced *in vivo* trophoblast differentiation potential compared to TSCs in teratoma assays. (A) Schematic view for the teratoma-forming assay. MEFs, TSCs and TELSCs were re-suspended and and subcutaneously injected into 6- to 8-week-old female NOG mice. (B) TSCs, TELSCs or control MEFs were injected subcutaneously into both flanks of NOG mice. Lesions were analyzed 10 days after injection. (C) Teratoma were dissected and fixed by H&E staining and IHC staining against the trophoblast marker ELF5 and endothelial marker CD31. The right image shows the morphological characteristics of TGCs with enlargement of nuclei. (D) IF staining of teratoma tissues derived from TELSCs and TSCs. STRA6: SynTI-specific marker. Scale bars, 200 μm. (E) Heatmap (left) and bubble chart (right) showing relative expression of the indicated genes in teratoma derived from MEFs, TSCs and TELSCs. Trophoblastic lineages data are from ^39^. (F) GSVA analysis of teratoma derived from MEFs, TSCs and TELSCs. (G) GSEA analysis of teratoma from MEFs, TSCs and TELSCs based on SynTI geneset. (H) UMAP visualization showing cells from TELSCs and TSCs teratomas colored by cell types. (I) Dot plots indicating the expression of cluster-specific genes in teratoma from TELSCs. (J) Pie chart indicating the proportion of cell types from TELSCs and TSCs teratomas. (K) Histogram indicating the ratio of SynTI from TELSCs and TSCs teratomas and ratio of original cell in teratomas derived from TELSCs and TSCs. (L) Heatmap showing DEGs among TELSCs, TSCs, TELSC-derived Ter-stem cells isolated from teratomas, and residual TSC-like cells (Ter-TSC) present in TSC-derived teratomas. Representative marker genes and enriched Gene Ontology (GO) terms are highlighted.

To precisely assess the cell fate commitments of TELSCs and TSCs, we performed qPCR, bulk RNA-seq, and scRNA-seq analyses. These analyses clearly showed that, compared with the control sample, a large number of trophoblast-specific, but not embryonic linage-specific, marker genes, were activated in both TSC- and TELSC-derived teratoma tissues, showing the TE developmental specificity of TELSCs (**Figures 4E, 4H, S4C and S4D**). Interestingly, bulk RNA-seq and qPCR analysis showed that the expression of a large group of mature trophoblast-specific genes, including *Crip1* (a marker for LaTP), *Serpinh1* (GlyT), *Glis1* (SynTI), *Col13a1* (SynTII), *Ghrh* (S-TGCs), *Flt1* (SpT), and *Prl2c2* (P-TGCs), was much higher in TELSC-derived teratomas than in the TSC-derived ones (**Figures 4E and S4E; Table S3**). Further GSEA and GSVA analyses consistently highlighted the superior trophoblast differentiation capacity of TELSCs, especially their unique ability to generate SynTI cells, a lineage not observed in TSC-derived teratomas (**Figures 4F, 4G; Table S4**), as further confirmed by immunofluorescence staining of teratomas (**Figure 4D**).

Precisely, scRNA-seq demonstrated that both TELSCs and TSCs gave rise to various trophoblast lineages, except for SpA-TGCs—known to require maternal signaling cues within the placenta (**Figures 4H, 4I, S4F and S4G**). While compared to TELSCs that can produce all other trophoblast lineages, TSCs failed to generate SynTI cells and rarely produced LaTP cells, which was quite consistent with RNA-seq and qPCR analysis (**Figures 4H, 4J, 4K and S4C**). Interestingly, we observed that a large group of undifferentiated TSCs, marked by *Rangrf*, *Nap1l1*, *Fbo21*, and *Efna3*, remained in TSC-derived teratoma tissues. In comparison, undifferentiated TELSCs were barely detectable in TELSC-derived ones, indicating that TELSCs have much higher differentiation potential than traditional TSCs (**Figures 4H and 4K**). This observation was quite consistent with qPCR and RNA-seq analysis, which showed significantly higher expression levels of stem cell marker genes in TSC-derived teratomas than in TELSC-derived ones (**Figures S4C and S4H**). In contrast, within TELSC-derived teratomas, we identified a unique group of ExE-like cells, marked by *Elf5*, *Hspd1*, *Krt17*, and *Amot*, accounting for approximately 4% of the total cell population, which could not be detected in TSC-derived tissues, and we named this population “Ter-stem” (**Figures 4H, 4I, 4J and S4F**). Interestingly, we detected 2,116 genes related to the TGF-beta and Notch signaling pathway, including *Tfap2a*, *Apoe, Smad3* and *Krt17*, that were specifically enriched in these Ter-stem cells, but not in TSCs. Comparatively, teratoma-retained undifferentiated TSCs were highly enriched with 1,660 genes related to Wnt signaling pathways, which were highly expressed in original TSCs, yet could not be detected in TELSC-derived Ter-stem cells in teratoma tissue (**Figure 4L**). Thus, TELSC-derived Ter-stem cells in teratoma tissue were distinct from known TSCs.

Collectively, in the teratoma assay, TELSCs displayed higher trophoblast developmental potency than TSCs and produced almost all mature trophoblast lineages and a unique population of ExE like Ter-stem cells distinct from TSCs.

### TELSCs can efficiently generate trophoblast organoids with mature trophoblast lineages and self-renewal ability

Next, we tried to assess the differentiation capacity of TELSCs *in vitro* and the related application in trophoblast organoid formation. We first constructed spontaneous differentiation by plating TELSCs in TX medium lacking FGF4, Heparin, and TGFβ1 for approximately 9 days, and we observed discernible morphological transformations toward trophoblast lineages (**Figures 5A and S5A**). qRT-PCR analysis revealed trophoblast lineage-specific genes, including *Syna* and *Plxnd1* for SynTI, *Synb* and *Gcm1* for SynTII, *Ascl2* and *Tpbpa* for SpT, and *Prl2c2* and *Ctsq* for TGC, underwent gradual and pronounced activation during the directed differentiation (**Figure S5B**). Further immunostaining analysis confirmed the appearance of corresponding lineages, such as SynTI (labeled by STRA6 and E-CADHERIN), IGF1R for SynTII, SpT (labeled by TPBPA and CDX2), and TGC (labeled by PROLIFERIN), after 9 days of induction (**Figure 5B**). Hence, it can be concluded that TELSCs possess the capability to generate diverse trophoblast lineages under the withdrawal of FGF4, Heparin, and TGFβ1 in two-dimensional (2D) culture conditions.

**Figure 5.**
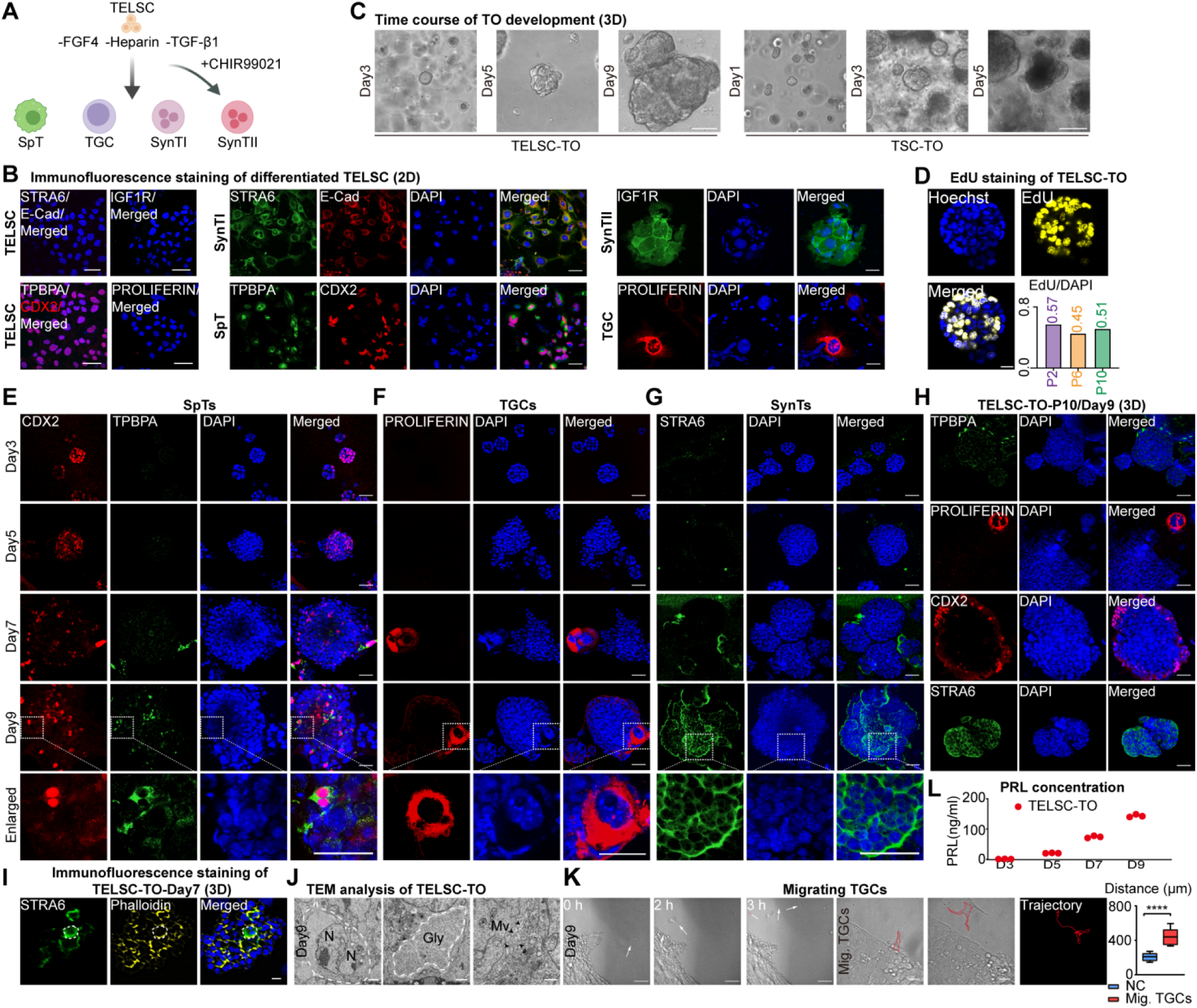
TELSCs, but not TSCs, can efficiently generate trophoblast organoids with mature trophoblast lineages and self-renewal ability. (A) Schematic of the procedures used to induce downstream differentiation of TELSCs in 2D culture condition. (B) Immunofluorescence staining of marker genes of downstream trophoblast cells (STRA6, IGF1R, TPBPA and PROLIFERIN) in differentiated TELSCs. Scale bars, 50 μm. (C) Morphology of trophoblast organoids derived from TELSCs and TSCs at different time points. Scale bars, 100 μm. (D) EdU staining of long-term cultured TELSC-TO. Scale bars, 25 μm. The ratio of EdU/DAPI is shown in the lower right. (E-G) Immunofluorescence staining of trophoblast organoids. TPBPA for SpT, PROLIFERIN for TGC and STRA6 for SynTI. Scale bars, 50 μm. (H) Immunofluorescence staining of trophoblast organoids in P10. TPBPA for SpT, PROLIFERIN for TGC and STRA6 for SynTI. Scale bars, 50 μm. (I) Immunofluorescence staining of trophoblast organoids showing the STRA6^+^ multinucleate cell. Scale bars, 20 μm. (J) Transmission electron microscope (TEM) view of TELSC-TO. Multinucleate cell, glycogen and microvillus were shown. N, nucleus; Gly, glycogen; Mv, microvillus. (K) The migration path of TGCs from TELSC-TO and quantification of the migration distance of TGCs. Scale bars, 25 μm. Mean ± SD; n = 10; ****p value < 0.0001, unpaired Student’s t test. (L) ELISA analysis of prolactin (PRL) in the medium secreted by TELSC-TO from day 3 to day 9 (n = 3).

The placenta, composed of diverse trophoblast lineages, plays a crucial role in fetal-maternal communication, yet the *in vitro* model mimicking the entire placenta development was still lacking. We then tested the possibility of deriving trophoblast organoids (TOs) from TELSCs by plating TELSCs, along with TSCs as the control, into modified human placental organoid culture medium, containing murine FGFβ, HGF, and EGF ^25–27^ for Matrigel-based 3D culture. After 3 days, we found the formation of organoid-like structures from TELSCs, which exhibited sustained and progressive growth over a period of at least 9 days (**Figure 5C**). In contrast, TSC-derived structures displayed impaired development with significant apoptosis after day 3 (**Figure 5C**). By day 9, TELSC-derived TOs developed into dense, solid masses. TUBULIN immunostaining, highlighting the cytoskeleton, confirms their well-organized, maturing structure (**Figure S5C**). Notably, these TELSC-derived TOs can be maintained through long-term passages (**Figure S5D**). While the EdU incorporation assay revealed the presence of proliferating cells across different passages, supporting the self-renewal capacity of TELSC-TOs (**Figures 5D and S5E**).

qRT-PCR analysis revealed a rapid decrease of TELSC-specific genes (*Cdx2*, *Elf5* and *Eomes*) after 3 days, indicating the initiation of differentiation. Correspondingly, trophoblast lineage markers, including *Syna* for SynTI*, Synb* and *Gcm1* for SynTII, *Ascl2* and *Tpbpa* for SpT, *Prl2c2* and *Ctsq* for TGC, exhibited dramatic induction after 7 days, signifying trophoblast specification (**Figure S5F**). While immunostaining analysis clearly showed that these TELSC-derived TOs, up to 10 passages, encompass various trophoblast lineages, such as SpT (TPBPA), TGC (PROLIFERIN), and SynTI (STRA6) (**Figures 5E-5I**). Using transmission electron microscopy (TEM), we detected the trophoblast cells with multiple nuclei sharing a continuous cytoplasm without intervening membranes, and the well-developed microvilli (Mv), representing SynT cells in TELSC-derived TOs. In addition, we observed glycogen granules (Gly) probably in the TGCs (**Figure 5J**). Notably, TGCs with large nuclei gathered together surrounding the trophoblast organoid and seemed able to migrate under living cell imaging (**Figure 5K**). Over 30% of these organoids also exhibited trophoblast outgrowth, mimicking placental invasion *in vivo* **(Figures S5G and S5H)**. It has been known that TGCs can produce prolactin (PRL), a hormone essential for pregnancy and the production of breast milk, and we then quantified PRL production in the placental organoids containing TGCs. We detected that TELSC-derived organoids gradually released increasing levels of PRL, reaching up to 150 ng/ml on the 9th day (**Figure 5L**). Thus, TELSCs can be widely used for trophoblast differentiation and TO generation formation.

### Newly identified ExE-like progenitors enable coupled self-renewal and differentiation abilities of TELSC-derived organoids

To uncover the comprehensive cell lineages within TELSC-derived TOs, we performed scRNA-seq analysis on day-9 trophoblast organoids derived from TELSCs and obtained approximately 5,088 single cells of high quality (**Figure 6A**). This analysis clearly showed that TELSC-derived organoids contained almost all reported differentiated trophoblast subpopulations (**Figures 6B and 6C**), including 640 SynTI cells (marked by *Batf3, Wnk3,* and *Tec),* 173 SynTII cells (marked by *Ank, Arhgap44, and Airn)*, 283 LaTPs (marked by *Cldn3, Cyba,* and *Adk)*, 300 SpTs (marked by *Maged1, Dapk2, and Tmem40*), 294 GlyTs (marked by *serpinb9e, Pitrm1, and Car2*), 634 S-TGCs (marked by *Hand1, Tmsb4x, and Bhlhe40*), and 300 P-TGCs (marked by *Prl3d1, Prl7a1, and Ctsl)* (**Figures 6A-6D and S6A; Table S5**).

**Figure 6.**
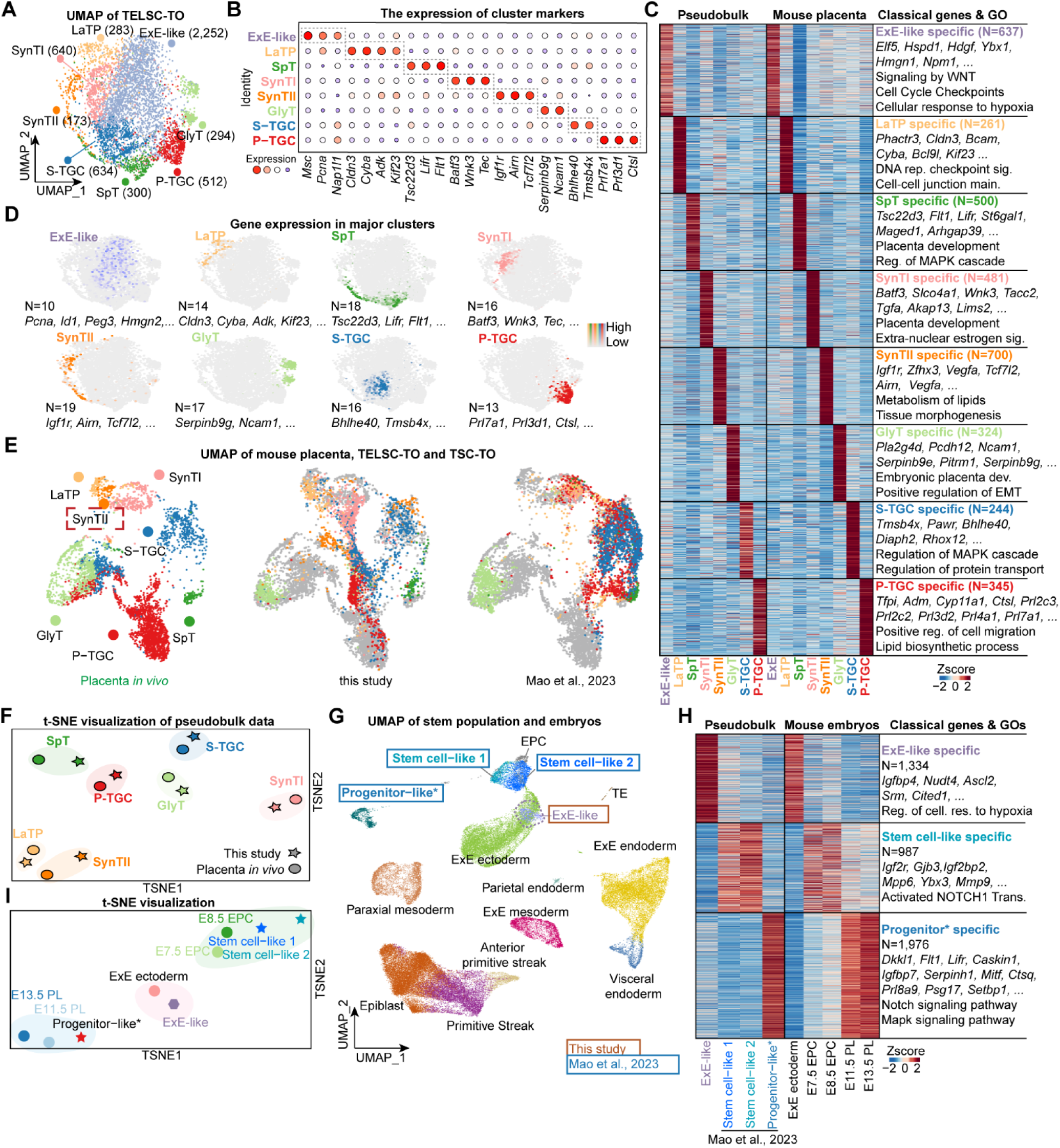
Newly identified ExE-like progenitors enable coupled self-renewal and differentiation abilities of TELSC-derived organoids. (A) Cellular composition of mouse TELSC-derived TO revealed by scRNA-seq. UMAP plot showing the 8 main clusters. (B) Dot plots indicating the expression of cluster-specific genes in TELSC-TO. (C) The heatmap displays the top differentially expressed genes across each identified cluster. Cluster-specific marker genes and representative Gene Ontology (GO) terms are listed on the right. (D) UMAP visualizations showing the expression of the marker genes in major clusters. (E) UMAP visualizations of the single-cell transcriptome of cells from TELSC-TO and the reported mouse trophoblast organoid integrated with mouse placenta. (F) t-SNE analysis of each cell types from TELSC-TO and mouse placenta. (G) Integration of stem cell population from TELSC-TO (ExE-like), TO reported by Mao et al., ^28^ and mouse embryo ^48^. (H) Heatmap on the left demonstrating the DEGs between ExE-like from TELSC-TO and stem cell population from ^28^ Heatmap on the right demonstrating the expression of each cluster in mouse embryos. (I) t-SNE analysis of stem cell population from TELSC-TO and the reported mouse trophoblast organoid.

In a recent study, Mao et al. claimed to generate TOs from TSCs, however, in which two different culturing media were required to maintain the proliferation of placenta organoids and to induce trophoblast differentiation, respectively ^38^. Thus these TSC-derived TOs are defective and distinct from TELSC-derived ones with coupled self-renewal and differentiation abilities that we developed in this study. To precisely compare the cellular compositions and understand the differences of these two TO types, we next performed integration analysis using scRNA-seq data from our TELSC-derived or published TSC-derived organoids, as well as the *in vivo* mouse placenta ^39^. This analysis clearly showed that TELSC-derived organoids contained almost all reported differentiated trophoblast subpopulations, which were highly comparable to the corresponding lineages in placenta tissue *in vivo*. Comparably, the TSC-derived TOs maintained in the trophoblast differentiation medium lacked the SynTII lineage (**Figures 6E and S6B**). Besides, the transcriptome-based t-SNE analysis clearly showed that various trophoblast cells in TELSC-derived TOs can be aligned well with those corresponding cell lineages in the placenta tissues *in vivo* (**Figure 6F**).

Interestingly, in the TELSC-derived organoids, we identified a unique, large population of 2,252 trophoblast progenitor cells, taking around 50% of all single cells obtained, which were enriched with typical trophoblast progenitor marker genes, such as *Fabp3*, *Igfbp4*, *Srm,* and *Pdgfa,* which were highly enriched in E5.5-6.5 ExE cells (**Figures 6A and 6D**). We therefore named these cells ExE-like progenitor cells. Further transcriptome-based clustering analysis clearly showed these ExE-like progenitor cells were indeed comparable to ExE ectoderm at E5.5-6.5, but not other stages (**Figure 6G**). Interestingly, stem cell-like and trophoblast progenitor-like populations were also identified in TSC-derived TOs cultured in the maintenance medium as recently reported. We therefore compared these TSC-derived stem cell-like populations, which were specially enriched with 987 genes, such as *Gjb3*, *Igf2bp2*, *Mpp6*, *Ybx3,* and *Mmp9,* which were particularly expressed in E7.5-8.5 ectoplacental cone (EPC) cells. Whereas, TSC-derived trophoblast progenitors particularly expressed 1,987 genes, including *Flt1*, *Lifr*, *Caskin1*, *Serpinh1*, *Prl8a9* and *Setbp1,* which were enriched in mature trophoblast lineages at around E11.5-13.5 stage. Comparably, there were 1,334 genes specially enriched in TELSC-derived ExE-like progenitor cells, which were highly expressed in E5.5-6.5 ExE ectoderm cells *in vivo* (**Figures 6H, S6C and S6D**). Further single-cell-based integration and t-SNE analysis clearly showed that TSC-derived stem cell-like and progenitor-like populations resembled E7.5-8.5 EPC cells and E11.5-13.5 trophoblast lineages, respectively. While TELSC-derived ExE-like progenitors were close to E5.5-6.5 ExE ectoderm cells *in vivo* (**Figure 6I**).

Finally, we noticed that ExE-like cells displayed elevated expression of proliferation-associated genes, suggesting a higher proliferative capacity and enhanced stemness compared to the stem cell-like and progenitor-like populations in TSC-derived TOs (**Figures S6E and S6F**). Summarily, TELSC-derived organoids with comprehensive mature trophoblast lineages and unique E5.5-6.5 ExE-like progenitors are clearly distinct from TSC-derived ones reported recently. These ExE-like cells with high proliferation capacity allow the coupled self-renewal and differentiation abilities of TELSC-derived organoids in a uniform medium for long-term passages.

### The rosette structure, relying on ITGB1, is required for ExE-like progenitor induction and TELSC-derived TO formation

Since these TELSC-derived ExE progenitor cells, TSCs, and recently reported TESCs all displayed ExE ectoderm features (**Figures 6G and 6H**), we then performed a transcriptome comparison on these cells. In brief, there were 712 genes that were specifically related to placenta development and cell growth, including *Peg10/3*, *Id3*, *Foxo4*, *Cdkn1c/11b* and *Mybl1*, highly expressed in the ExE-like cells and E5.5-6.5 ExE ectoderm cells, but not in reported TSCs or TESCs (**Figure 7A**). Additionally, we detected 974 genes related to stem cell proliferation (exemplified by *Car2*, *H19*, *Pou3f1,* and *Wnt3*) and 1356 genes associated with *in utero* embryonic development (such as *Pgk1*, *Apoe,* and *Apoc1*), all of which were specifically expressed in E5.6-6.5 ExE ectoderm cells and were specifically enriched in TESCs and TSCs, respectively. There were also 310 ExE-specific genes related to reproductive structure development, including *Fgf2*, *Sox4*, *Cebpb*, and *Nop10*, which were particularly highly expressed in both ExE-like progenitors and TESCs, but not TSCs (**Figure 7A and S7A**). Thus, ExE-like cells exhibited novel E5.5-6.5 ExE features distinct from conventional TSCs or newly reported TESCs (**Figure 7B and S7B**).

**Figure 7.**
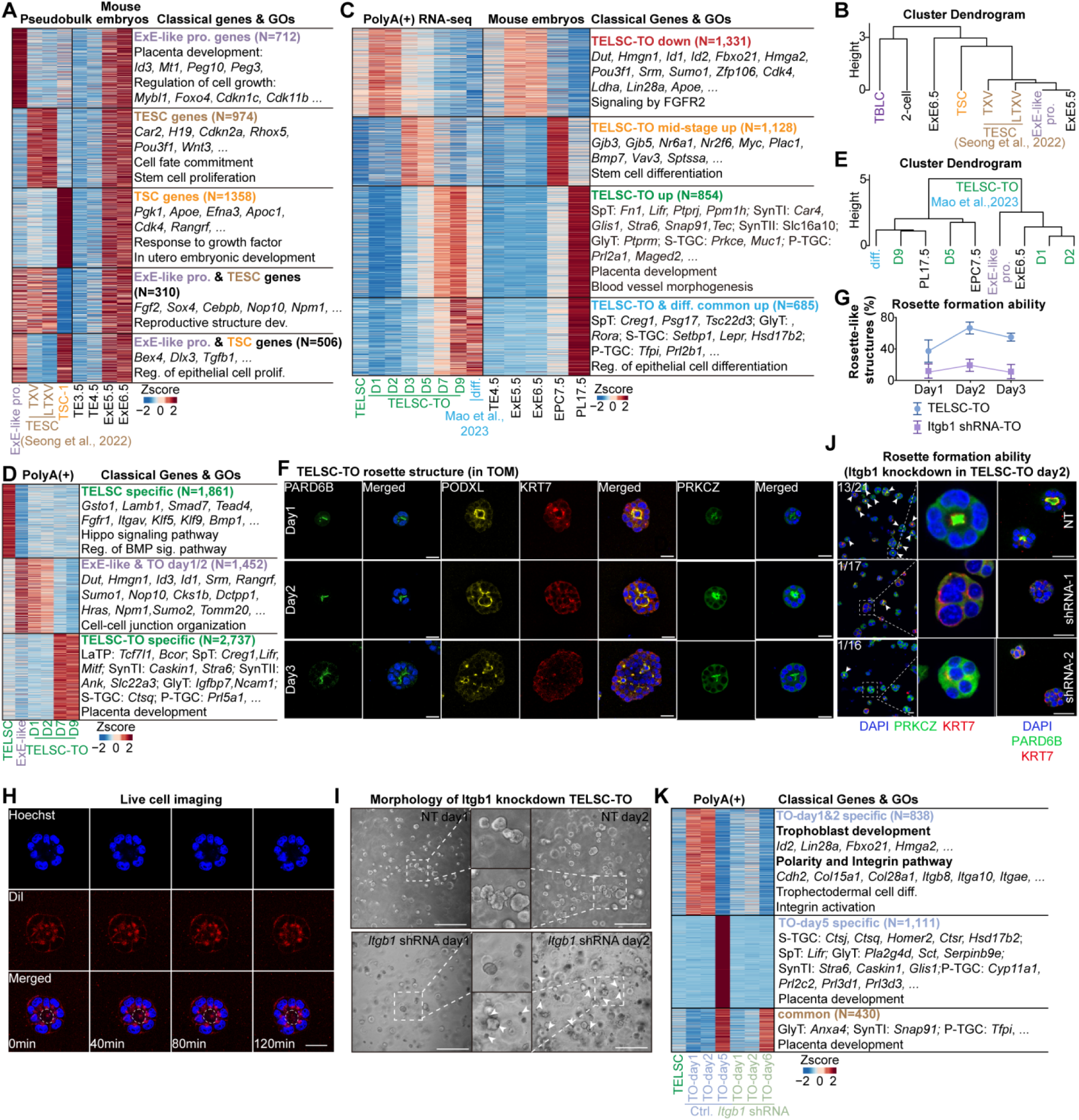
The rosette structure, relying on ITGB1, is required for ExE-like progenitor induction and TELSC-derived TO formation. (A) Heatmap on the left demonstrating the DEGs between stem cell population in TELSC-TO (ExE-like), TESCs and TSCs. Heatmap on the right demonstrating the expression of each cluster in mouse embryos. (B) Hierarchical clustering analysis of ExE-like, TESCs, TSCs, TBLCs and mouse embryos data. (C) Heatmap indicating the differentially expressed genes of TELSC-TO, TSC-TO, TO reported by Mao et al.,^28^ and mouse embryo ^44,45,49,50^. The classical genes and enrichment of GO terms of these genes is shown on the right. (D) The heatmap illustrates distinct transcriptional signatures among TELSCs, ExE-like cells identified in single-cell RNA-seq of TELSC-TOs, and bulk RNA-seq profiles of TELSC-TOs. Cluster-specific marker genes and representative GO terms are listed on the right. (E) Hierarchical clustering was performed based on transcriptomic profiles from the ExE-like population identified in TELSC-derived TOs (both single-cell and bulk RNA-seq), mTOs reported by ^28^, and corresponding *in vivo* mouse embryos. (F) Immunofluorescence staining of PARD6B, PODXL, KRT7, and PRKCZ in TELSC-TO from day 1 to day 3. Scale bars, 25 μm. (G) Line graph showing the percentage of TELSC-derived organoids that formed rosette-like structures at days 1, 2, and 3. (H) Live-cell imaging of rosette-like structures in TELSC-TO. Scale bars, 25 μm. (I) Morphology of TELSC-TO after transfected with NT and Itgb1 shRNA. (J) NT (non-targeting) and *Itgb1* shRNA-1 to −2 were transfected into TELSCs. The location of PRKCZ and PARD6B in TELSC-TO were measured by immunofluorescence staining. Scale bars, 25 μm. Quantification of the number of TELSC-TO with rosette-like structures transfected with NT and *Itgb1* shRNA-1 to −2 were labeled. (K) Heatmap indicating the differentially expressed genes of TELSCs, TELSC-TO and TELSC-TO transfected with *Itgb1* shRNA. The representative genes and enrichment of GO terms of these genes is shown.

To further precisely understand the induction of ExE-like progenitors during TELSC-derived organoid formation, we performed RNA-seq on TELSC-derived organoids at various time points, including days 1, 2, 3, 5, 7, and 9. More than 800 genes were specifically activated after 7-9 days in TELSC-based organoid formation, and these genes represented various trophoblast lineage-specific genes, including *Fn1* and *Mitf* for SpT, *Stra6* and *Car4* for SynTI, *Atxn1* and *Col13a1* for SynTII, *Col4a1*, and *Ptprm* for GlyT, *Arhgef25* and *Prkce* for S-TGC (**Figure 7C; Table S6**), indicating eventual commitment to distinct mature trophoblast lineages. In addition, approximately 936 genes were specifically activated between days 3 and 5, including *Gjb3* and *Mpp6*, which are primarily expressed during the embryonic ectoplacental cone (EPC) stage *in vivo*. Interestingly, a cluster of 645 genes was dynamically induced after 1-2 days but decreased rapidly after 3 days and was observed specifically during TELSC differentiation. Notably, these genes were associated with trophoblast progenitors, including *Dut*, *Id3*, *Hmgn1,* and *Id1,* which were also highly expressed in E5.5-6.5 ExE ectoderm, and ExE-like progenitors identified in TELSC-derived TOs (**Figure 7D**). Further clustering analysis and GSEA indicated that these day 1/2 cells closely resembled ExE-like progenitors in TELSC-derived TOs; day 5 cells displayed transcriptomic features akin to those of the epc stage; whereas, in comparison, cells from day 7-9 organoids showed transcriptomic profiles similar to those of mature placental tissues at E9.5 and E17.5 (**Figure 7E and S7C**). Thus, ExE-like progenitors, representing an intermediate phase for trophoblast progenitor expansion, were induced at days 1-2 during TELSC differentiation.

During *in vivo* mouse embryo development at around E5.5-6.5, both embryonic epiblast and extraembryonic ExE cells form a rosette-like structure with apical domains, showing an unpolarized-to-polarized transition, which is essential for lumenogenesis of developing embryos and the related cell differentiation process ^12,40,41^. Interestingly, immunostaining analysis using antibodies against PARD6B and PODXL, well-known apical domain markers, clearly showed that typical rosette-like structures with the expression of PARD6B and PODXL at the lumen center were transiently and dynamically induced on days 1 and 2, yet disappeared after day3, during TELSC-derived organoid formation, regardless of whether our own culture system or that described by Mao et al^28^ was used (**Figures 7F,7G and S7D**). Further live cell imaging using Dil staining at 0, 40, 80, and 120 minutes showed the rosette structure forms and maintains stability for a long time, indicating a stable cellular arrangement event (**Figure 7H**). In contrast, TSC-derived organoids did not efficiently generate rosette structures like TELSCs under both culture conditions (**Figures S7E and S7F**), showing the functional deficiency of TSCs on embryo-like structures compared to TELSCs. Since the induction of the above rosette-like structures and ExE-like progenitors occurred at the same time, we proposed that the rosette-like structure formation was essential for ExE-like progenitor cell fate determination.

To test the above hypothesis, using shRNAs, we knocked down the expression of *Itgb1*, a key integrin β1 signaling gene required for apical domain formation, in TELSCs and subsequently assessed their capacity to form rosette structures. qPCR was performed, and confirmed the efficient knockdown of *Itgb1*, which interestingly also significantly reduced mRNA levels of polarity markers, including *Pard6b*, *Podxl*, and *Prkcz,* in the cells after 2 days of induction for TO formation (**Figure S7G**). Immunofluorescence staining further revealed a significant reduction in the proportion of rosette structures (**Figure 7I**), eventually leading to peripheral cell death beginning after 2 days (**Figure 7J**). RNA-seq was then performed on organoid samples from day 1 to day 6 post-shRNA treatment. As expected, we found ITGB1 knockdown obviously inhibited the dynamic activation of 838 genes, involved in extracellular matrix organization (e.g., *Col15a1*, *Col28a1*), cell adhesion (e.g., *Itgb8*, *Itga10*), and ExE-stage identity (e.g., *Id2*, *Lin28a,* and *Fbxo21*), which were highly enriched in ExE-like progenitors in TELSC-derived organoids and ExE cells *in vivo*, therefore inhibiting the induction of ExE-like cells at days 1 and 2 (**Figure 7K**). Consistently, GSEA analysis confirmed significant downregulation of pathways related to integrin signaling and cell polarity following ITGB1 knockdown (**Figure S7H**). At later stages, these knockdown organoids failed to fully develop. RNA-seq analysis of day 6 samples revealed reduced expression of multiple lineage-specific markers compared to the WT group, including *Ctsj* (S-TGC), *Lifr* (SpT), and *Pla2g4d* (GlyT), indicating that early disruption of rosette morphogenesis substantially impairs subsequent lineage specification (**Figure 7K**).

Altogether, using the TELSC-derived organoid model, we demonstrate that the formation of rosette-like structures, relying on the key integrin β1 signaling factor *Itgb1*, is indispensable for ExE-like progenitor induction and further TO generation from TELSCs, which could explain the deficiency of TSCs in TO formation. Nevertheless, capturing TELSCs enables us to faithfully recapitulate key morphogenetic events of the entire trophoblast development *in vitro*, showing the widespread applications in basic studies and translational medicine.

## Discussion

In our recent study, we have successfully established the capture and long-term maintenance of human and mouse totipotent stem cells, TBLCs, comparable to 2- and 4-cell stage blastomeres, through spliceosomal repression^29,42^. Our further investigation into the differentiation potential of these cells toward various functional cells, particularly the extraembryonic lineages containing trophoblasts, highlights the unique developmental potency of TBLCs. Here, based on the Hippo-YAP/Notch-to-TGFβ1 signaling switch, we developed a “two-step” differentiation system to robustly and efficiently induce TBLCs or 8-cell blastomeres to produce a new kind of trophectoderm-like stem cells (TELSCs), reassembling the E4.5 TE cells, which are distinct from the well-known TSCs or newly reported TESCs ^24^ and TSCs ^21^. Importantly, TELSCs exhibit specific trophectoderm properties in both teratoma differentiation and chimera formation assays. We demonstrated that TELSCs can broadly produce all major placental trophoblast lineages, including LaTPs, GlyTs, S-TGCs, SpA-TGCs, P-TGCs, SpTs, SynTIs, and SynTIIs, at the single-cell level for the very first time. Therefore, our study provides a very useful platform and model to study the earliest events of trophoblast development from totipotent cells.

In our 3D culture system, TELSCs resembling TE4.5 transit through an E5.6-E6.5 ExE-like phase before differentiating into mature trophoblast organoids with diverse lineages. Interestingly, transcriptomic comparison showed that these ExE-like progenitors were clearly different from reported TSCs or TESCs, although both of which have partial features of E5.5-6.5 ExE ectoderm *in vivo*. Notably, the production of these ExE-like progenitors highly relies on the morphological change with the formation of rosette-like structures, which were governed by the key integrin β1 signaling gene, *Itgb1.* These progenitors with rapid cell proliferation capacity can be persistently retained in TELSC-derived organoids, which therefore enabled the formation of TOs with coupled self-renewal and differentiation abilities for long-term passages. Comparably, TSCs cannot undergo similar rosette structure transformation, and therefore cannot generate organoids in the same culturing condition, which might reflect the deficiency of conventional TSCs with post-implantation characters compared to TELSCs we captured in this study. In contrast, two types of culturing medium were newly developed and required to maintain the proliferation and differentiation of TSC-derived TOs that failed to produce mature SynTII cells respectively.

Our newly established mouse trophoblast organoids from TBLCs offer a comprehensive model for studying mouse placental development, facilitating high-throughput genetic screening and in-depth investigation into pre-, peri-, and post-implantation processes. This is pivotal for exploring the pathophysiological mechanisms underlying implantation failures and pregnancy disorders, such as miscarriage and preeclampsia, which are frequently attributed to defective placentation. This *in vitro* system provides critical insights into these reproductive disorders, advancing our comprehension of trophoblast differentiation from totipotent stem cells and providing a powerful platform for placental development research.

### Limitations of study

While mouse TELSCs can be efficiently derived from TBLCs to form placental trophoblast organoids, the application of this system in basic research and reproductive medicine should be widely explored. Additionally, the co-culture system containing the TELSC-derived trophoblast organoid, maternal decidual cells, as well as the immune and endothelial cells needs to be further developed, which will be very important to fully understand the fetal-maternal communications during the establishment and maintenance of pregnancy. Finally, whether the similar culture conditions can be applied to capture human TELSC and TELSC-derived organoids could be a very interesting research topic.

## Acknowledgments

We thank the Flow Cytometry Core at National Center for Protein Sciences at Peking University for technical help and appreciate the Core Facilities of Life Sciences, Peking University, for the assistance with confocal microscopy and 10x Genomics SC Chromium. We are grateful to the Imaging Core Facility, the State Key Laboratory of Membrane Biology. This work was supported by grants to P.D. from the National Key Research and Development Program of China (2024YFA1106900 and 2021YFA1100100), the Beijing Outstanding Young Scientist Program (JWZQ20240101003), and the Beijing Natural Science Foundation (Z230011), the National Natural Science Foundation of China (32225017); and a postdoctoral fellowship to B.P. from the China Postdoctoral Science Foundation (2025T180736), as well as partial support from the Yifu Outstanding Postdoctoral Award.

## Author Contributions

X.J., B.P., Z.H., and C.W. performed most of the experiments, X.J. and C.W. performed all the bioinformatic analyses. P.D., and W.T. designed all experiments and analyzed data. P.D. conceived and supervised this project and wrote the paper with and B.P.

---

**Figure S1.**
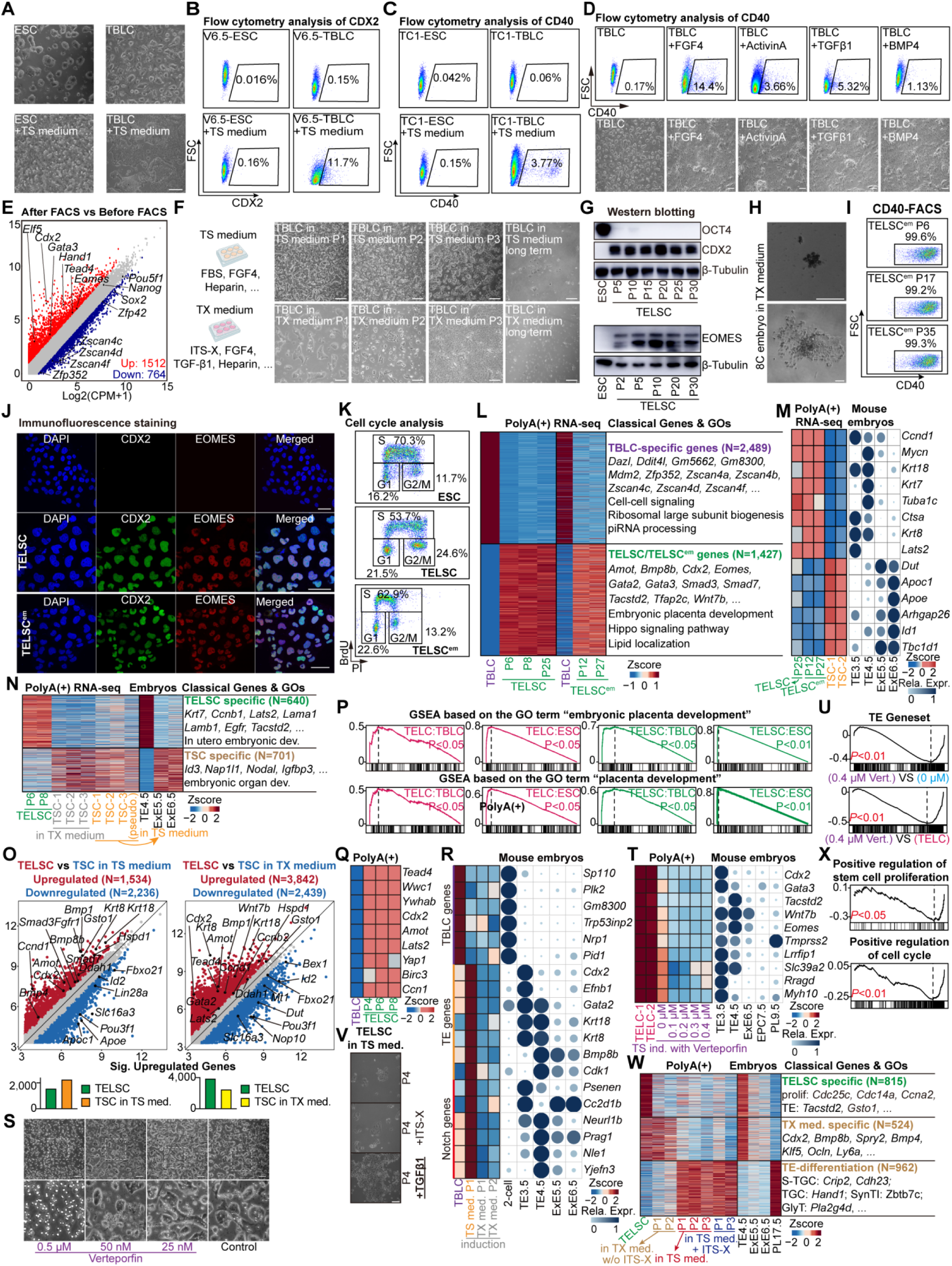
Capturing trophectoderm-like stem cells (TELSCs) with pre-implantation E4.5 TE features using a “two-step” culture system, related to Figure 1. (A) The morphology of ESCs, TBLCs and ESCs, TBLCs in TS medium after 3 days of induction. Scale bars, 250 μm. (B) FACS analysis of the percentage of CDX2^+^ cells from ESCs and TBLCs, as well as ESCs and TBLCs cultured in TS medium, using V6.5 cell line. (C) FACS analysis of the percentage of CD40^+^ cells from ESCs and TBLCs, as well as ESCs and TBLCs cultured in TS medium, using TC1 cell line. (D) FACS analysis of the percentage of CD40^+^ TELCs obtained from the TBLCs after induction with different molecules, including FGF4, Activin A, TGFβ1 and BMP4. The corresponding cell morphology is displayed in the lower panel. (E) Scatterplots displaying the transcriptome comparison of TELCs before and after CD40-based FACS using RNA-seq. Upregulated (FC>2) and downregulated (FC<0.5) genes are shown in red and blue, respectively. (F) The morphology of TBLCs of different passages and long-term culture in TX and TS medium, also the morphology of TBLCs after CD40 FACS after induction. Scale bars, 250 μm. (G) Western blotting was used to detect OCT4, CDX2 and EOMES in TELSCs from different passages. β-Tubulin was used as a loading control. (H) The morphology 8C embryos cultured in TX medium. Scale bars, 250 μm. (I) FACS analysis of the percentage of CD40^+^ cells in TELSC^em^s at different passages. (J) Immunofluorescence staining of TFAP2C and PEG10 in TBLCs, TELSCs and TELSC^em^s. Scale bars, 50 μm. (K) Cell cycle analysis of ESCs, TELSCs and TELSC^em^s. (L) Heatmap indicating the relative expression of TBLCs, TELSCs and TELSC^em^s. The representative genes and enrichment of GO terms of these genes is shown. (M) Heatmap indicating the relative expression of characteristic genes in TELSCs, TELSC^em^s and TSCs. Bubble chart showing the relative expression of these genes in mouse embryos. (N) Heatmap indicating the relative expression of characteristic genes in TELSCs, TSCs cultured in TX medium and TSCs cultured in TS medium. Heatmap on the right demonstrating the expression of each cluster in mouse embryos. The representative genes and enrichment of GO terms of these genes is shown. (O) The scatter plot displays differentially expressed genes between TELSCs and TSCs cultured in various media. The bar graph summarizes the number of differentially expressed genes identified under each comparison condition. (P) GSEA analysis of ESCs, TBLCs, TELCs and TELSCs based on “embryonic placenta development” and “placenta development” geneset. (Q) Heatmap indicating the differentially expressed genes in Hippo pathway of TELSCs and TBLCs. (R) Heatmap indicating the relative expression of characteristic genes in TELSCs, TSCs cultured in TX medium and TSCs cultured in TS medium. Bubble chart showing the relative expression of these genes in mouse embryos. (S) Phase contrast images of TBLCs cultured in TS medium for 24h supplemented with Verteporfin at the indicated concentration. Scale bars, 100 µm. (T) Heatmap indicating the differentially expressed genes of TELCs and TBLCs induction in TS medium plus verteporfin. Bubble chart showing the relative expression of these genes in mouse embryos. (U) GSEA analysis of TELCs, TBLCs induction in TS medium and in TS medium plus verteporfin based on TE geneset. (V) The morphology of TELSCs cultured in TS medium, TS medium plus ITS-X and TS medium plus TGFβ1. (W) Heatmap indicating the differentially expressed genes of TELSCs, TBLCs induction in TX medium withdraw ITS-X, in TS medium and in TS medium plus ITS-X. Heatmap on the right demonstrating the expression of each cluster in mouse embryos. The representative genes and enrichment of GO terms of these genes is shown. (X) GSEA analysis of TBLCs induction in TX medium withdraw ITS-X and in TX medium based on “Positive regulation of stem cell proliferation” and “Positive regulation of cell cycle” geneset.

**Figure S2.**
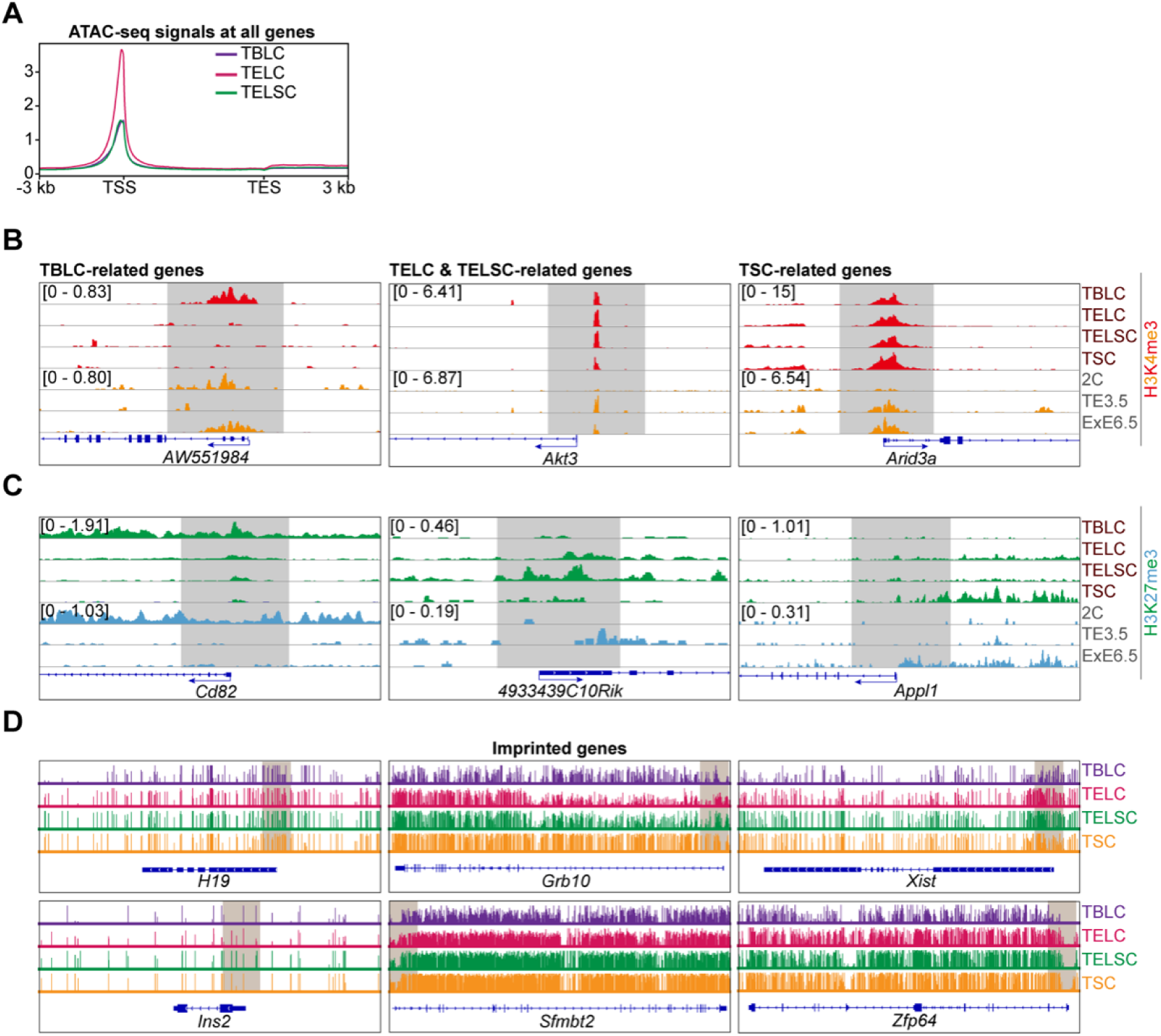
The unique epigenomic features of TELCs and TELSCs, related to Figure 2. (A) Average ATAC-seq signals at all genes in TBLCs, TELCs and TELSCs. (B) IGV browser view displaying H3K4me3 signals of specific genes in TBLCs, TELCs, TELSCs and TSCs, and mouse embryos ^35,36^. (C) IGV browser view displaying H3K27me3 signals of specific genes in TBLCs, TELCs, TELSCs and TSCs, and mouse embryos ^35,36^. (D) IGV browser view displaying DNA methylation patterns of imprinting genes in TBLCs, TELCs, TELSCs and TSCs.

**Figure S3.**
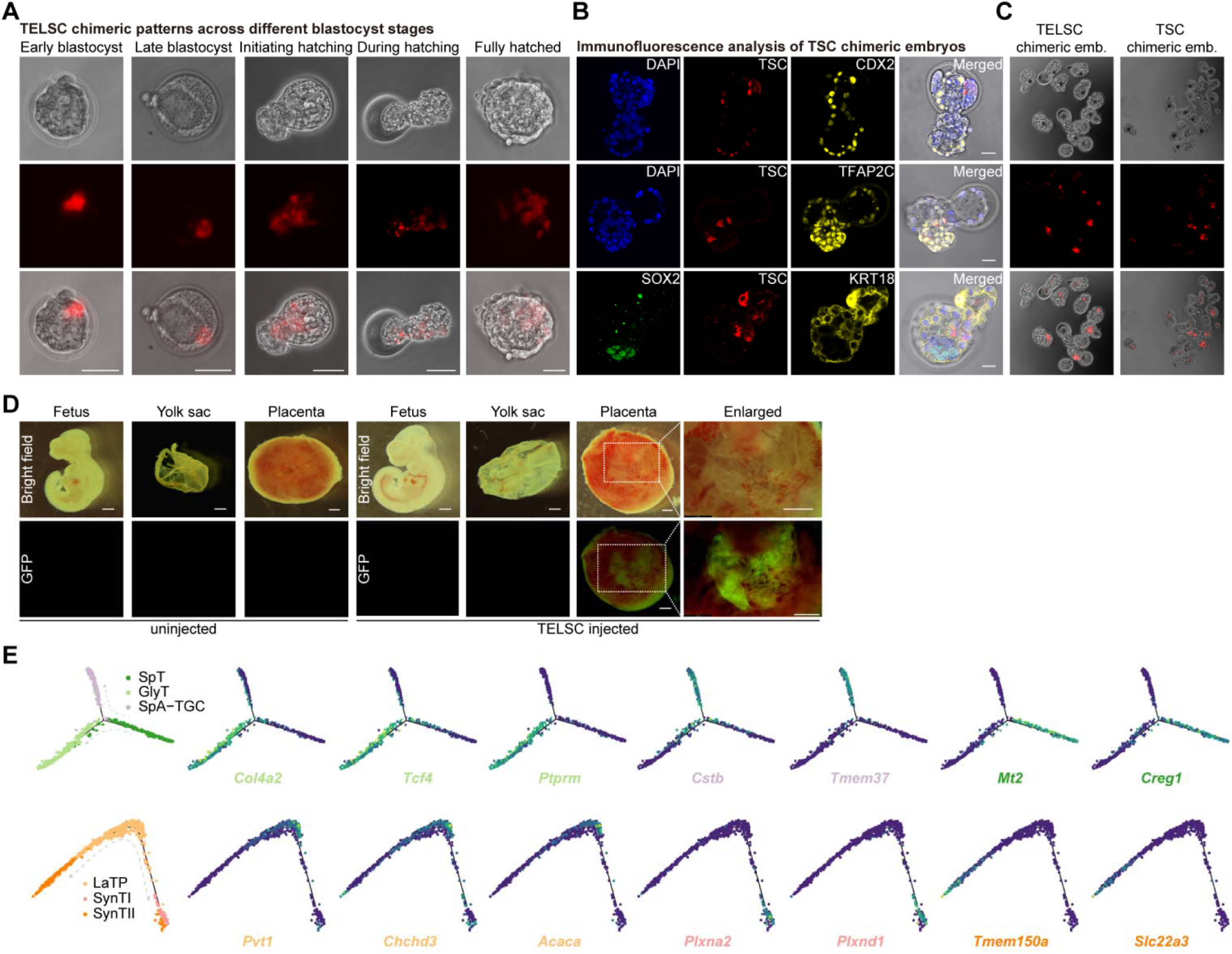
TELSCs exhibit robust *in vivo* developmental potential and full trophoblast lineage contribution, related to Figure 3. (A) Representative images of chimeric embryos injected mCherry-labeled TELSCs. TELSCs can cooperate into trophectoderm across blastocyst developmental stages: early blastocyst, late blastocyst, hatching-initiation blastocyst, hatching-progress blastocyst, and fully hatched (zona pellucida-free) blastocyst. Scale bar, 50 μm. (B) Representative immunostaining of chimeric blastocysts injected with mCherry-labeled TSCs. CDX2, TFAP2C and KRT18: TE-specific markers; SOX2: ICM-specific marker. Scale bar, 20 µm. (C) Images of chimeric embryos injected with mCherry-labeled TELSCs or TSCs. Asterisk, TELSCs or TSCs contribute to TE and form chimeric embryos. (D) Images of chimeric conceptuses derived from 8-cell embryos injected with donor EGFP-labeled TELSCs with an uninjected conceptus as control. Scale bar, 1 mm. (E) Monocle2 analysis of lineage-specific gene expression of TELSC-chimeric placenta across single-cell differentiation trajectories.

**Figure S4.**
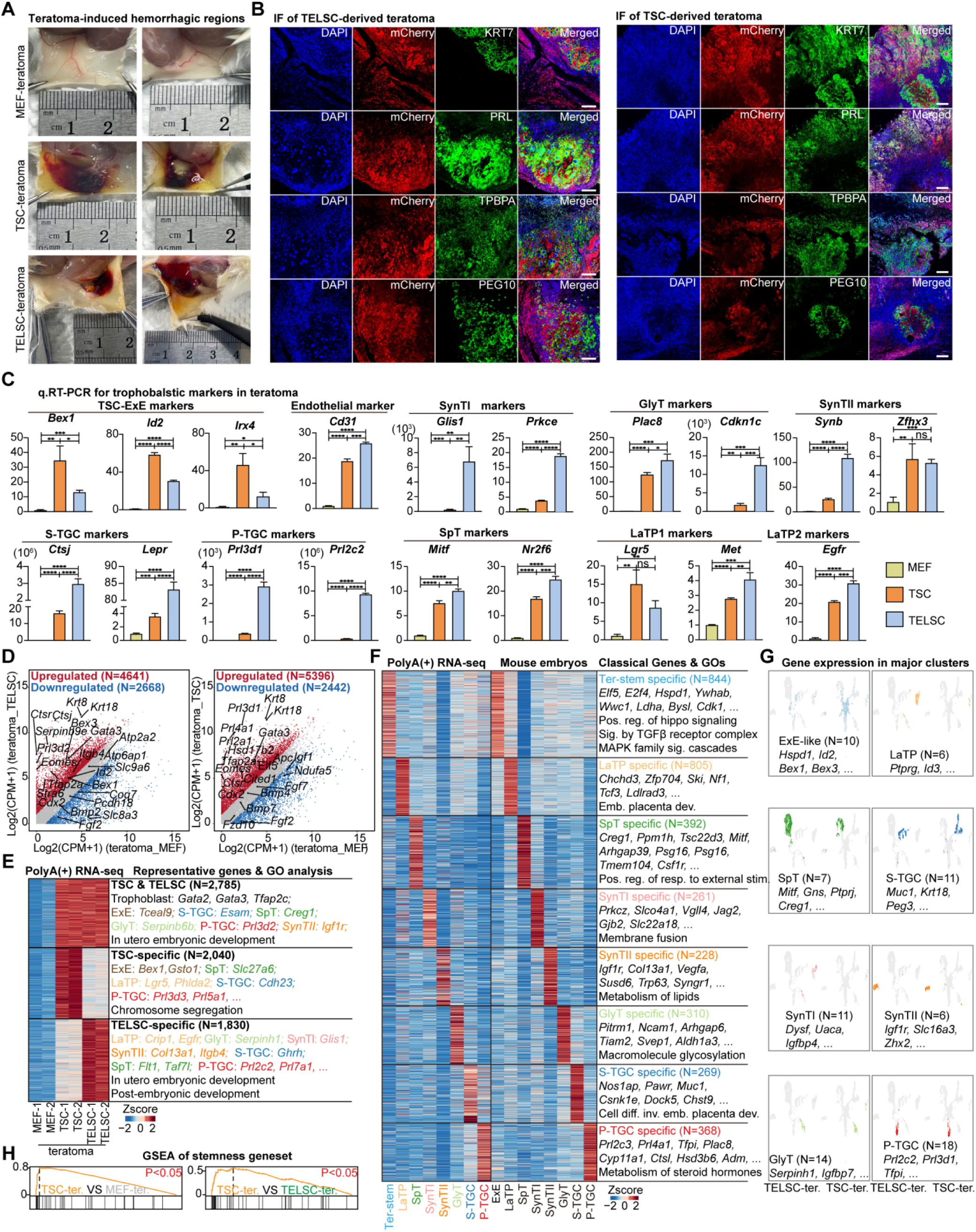
TELSCs exhibit enhanced *in vivo* trophoblast differentiation potential compared to TSCs in teratoma assays, related to Figure 4. (A) TSCs, TELSCs or control MEFs were injected subcutaneously into both flanks of NOG mice. Lesions were analyzed 10 days after injection. Replication of (Figure 4B), demonstrating the reproducibility of the results. (B) Immunofluorescence analysis of teratoma tissues derived from TELSCs and TSCs. Scale bars, 200 mm. (C) qRT-PCR analysis of the relative expression of the trophoblast lineage marker of teratoma derived from MEFs, TSCs and TELSCs. Data were normalized to GAPDH. Mean ± SD; n = 3 technical replicates and statistically significant differences (Student’s t test) are indicated with asterisks. (D) Scatterplots displaying the transcriptome of teratoma derived from MEFs, TSCs and TELSCs using RNA-seq. Upregulated (FC > 2) and downregulated (FC < 0.5) genes are shown in red and blue, respectively. (E) Heatmaps of the relative expression of commonly and specifically upregulated genes in teratoma derived from TSCs, TELSCs compared with teratoma derived from MEFs. The representative genes and enrichment of GO terms of these genes is shown. (F) The heatmap displays differentially expressed genes across clusters identified in TELSC-derived teratomas. The expression profiles of these genes are also shown in corresponding stages of *in vivo* mouse placental development. (G) UMAP visualizations showing the expression of the marker genes in major clusters in teratoma derived from TSCs, TELSCs. (H) GSEA analysis of teratoma derived from MEFs, TSCs and TELSCs based on stemness geneset.

**Figure S5.**
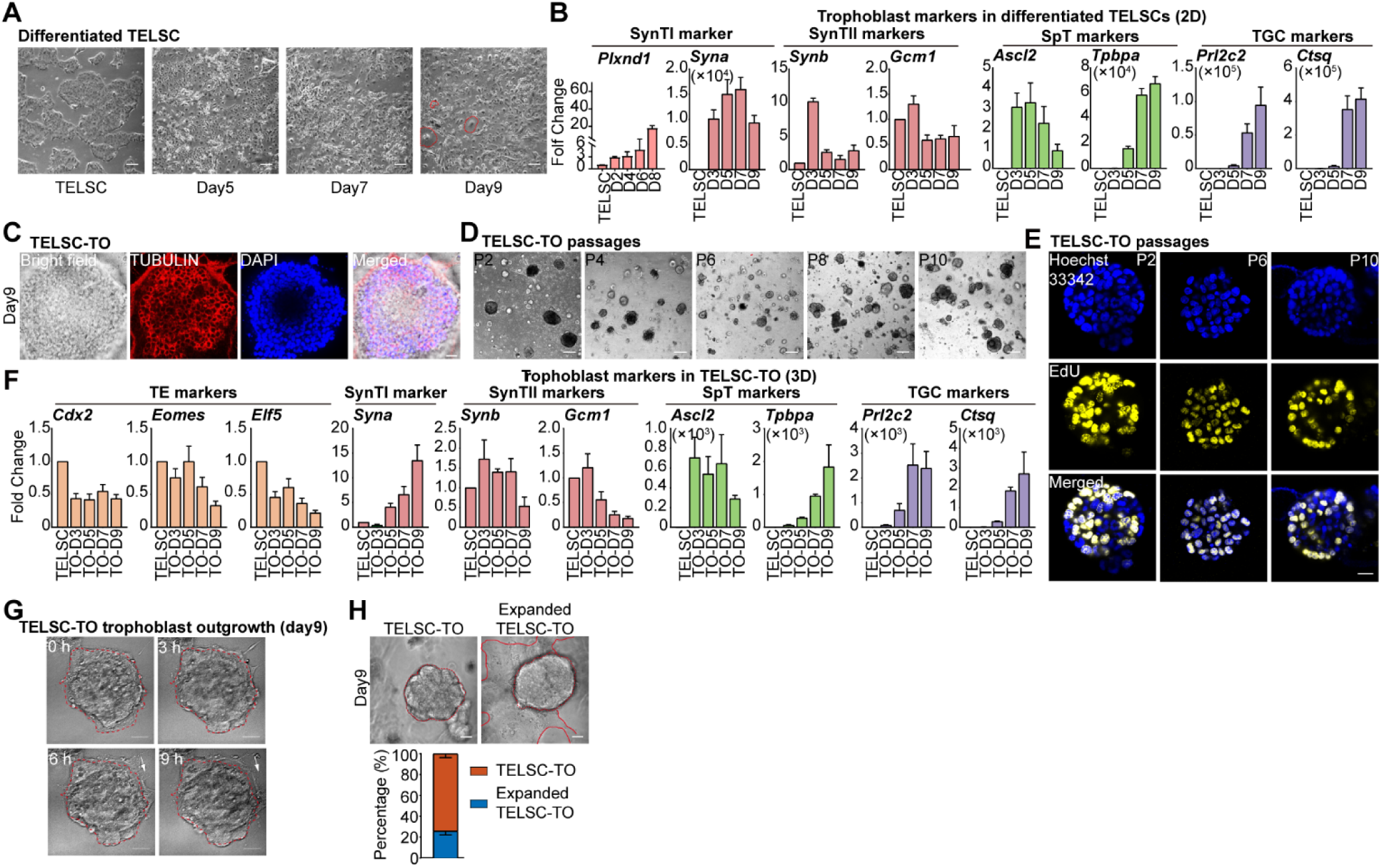
TELSCs, but not TSCs, can efficiently generate trophoblast organoids with mature trophoblast lineages and self-renewal ability, related to Figure 5. (A) Morphology of long-term cultured TELSCs and differentiated TELSCs in 2D culture condition. Scale bars, 25 μm. (B) qRT-PCR analysis of the relative expression of different trophoblast markers in TELSCs and differentiated TELSCs in 2D culture condition at different time points. Data were normalized to GAPDH. Mean ± SD; n = 3 technical replicates. (C) The structure of TELSC-TO at day 9. Immunofluorescence staining of TUBULIN to show the cytoskeletal structure of TELSC-TO. Scale bars, 25 μm. (D) Morphology of long-term cultured TELSC-TO. Scale bars, 25 μm. (E) EdU staining of long-term cultured TELSC-TO at different passages. Scale bars, 25 μm. (F) qRT-PCR analysis of the relative expression of different trophoblast markers during TELSC-TO formation: *Cdx2*, *Eomes* and *Elf5* for TE; *Syna* for SynTI; *Synb* and *Gcm1* for SynTII; *Ascl2* and *Tpbpa* for SpT; *Prl2c2* and *Ctsq* for TGC. Data were normalized to GAPDH. Mean ± SD; n = 3 technical replicates. (G) Long-term morphological changes in TELSC-TO. The arrowheads represent the outgrowth of TELSC-TO. Scale bars, 25 μm. (H) Morphology of expanded TELSC-TO at day 9 and corresponding statistics. Scale bars, 25 μm.

**Figure S6.**
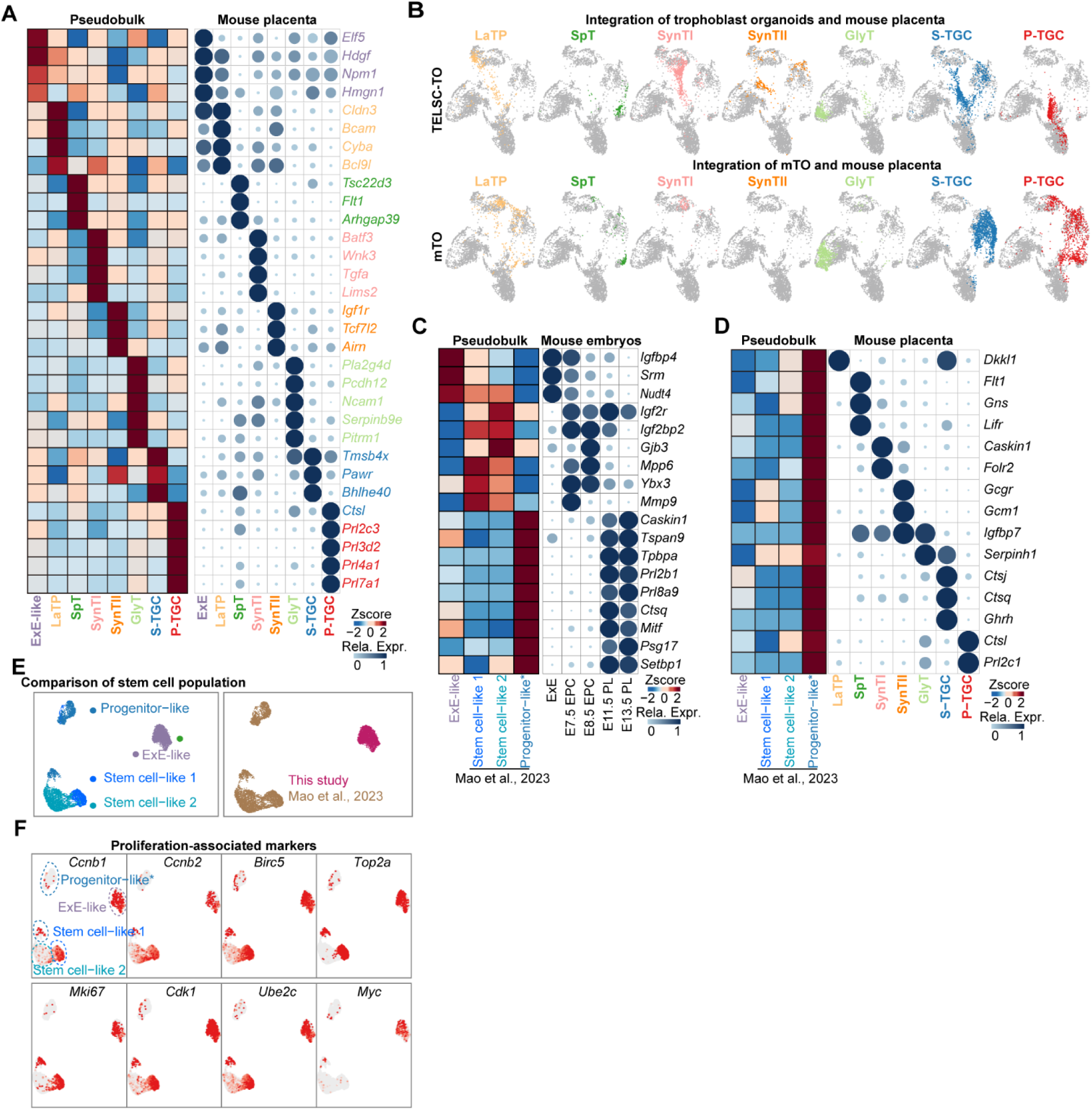
Newly identified ExE-like progenitors enable coupled self-renewal and differentiation abilities of TELSC-derived organoids, related to Figure 6. (A) Heatmap and bubble chart displaying marker gene expression in trophoblast cell types in TELSC-TO and mouse embryos. (B) UMAP visualizations showing the integration of TELSC-TO, TO reported by ^28^ and mouse placenta, the gray points represent mouse embryonic data. (C) Heatmap on the left demonstrating the DEGs between ExE-like from TELSC-TO and stem cell population from ^28^ Bubble chart showing the relative expression of these genes in mouse embryos. (D) Heatmap and bubble chart displaying marker gene expression in ExE-like, stem cell population from ^28^ and the reported mouse trophoblast cell types. (E) The integration of the scRNA-seq results of stem cell population from TELSC-TO (ExE-like) and from TO reported by ^28^. (F) UMAP visualizations showing the expression of the proliferation-associated marker genes in stem cell population.

**Figure S7.**
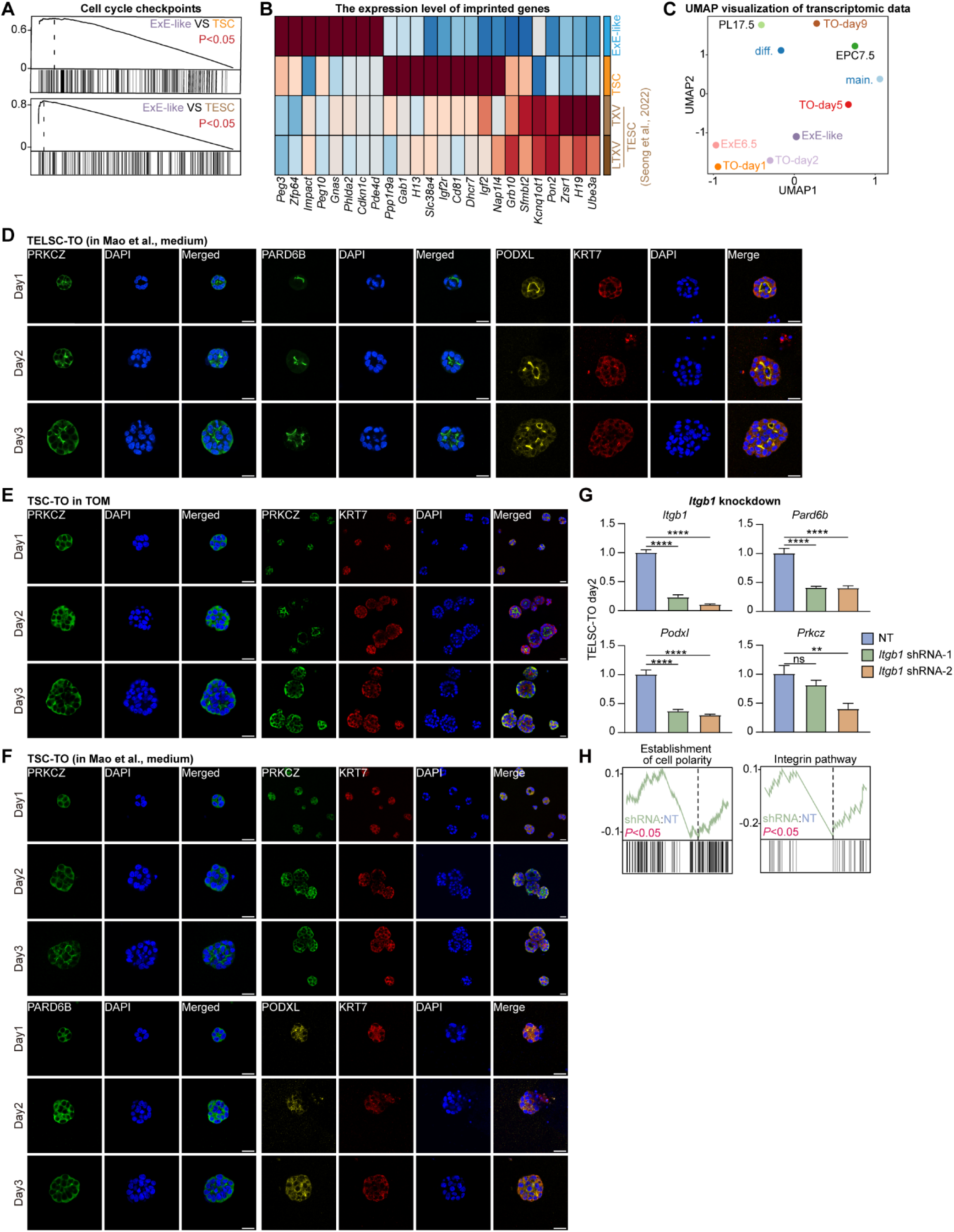
The rosette structure, relying on ITGB1, is required for ExE-like progenitor induction and TELSC-derived TO formation, related to Figure 7. (A) GSEA analysis of ExE-like, TSCs and TESCs based on “Cell cycle checkpoints” geneset. (B) Heatmap comparing placenta-associated imprinted gene expression across ExE-like, TSCs, and TESCs. (C) UMAP analysis of bulk RNA-seq of TELSC-TO, pseudo-bulk of mTO from ^28^ and also mouse embryos. (D) Immunofluorescence staining of PARD6B, PODXL, PRKCZ and KRT7 in TELSC-TO from day 1 to day 3. Scale bars, 25 μm. (E-F) Immunofluorescence staining of PARD6B, PODXL, PRKCZ and KRT7 in TSC-TO from day 1 to day 3. Scale bars, 25 μm. (G) qRT-PCR analysis of the relative expression of *Itgb1*, *Pard6b*, *Podxl* and *Prkcz* after transfected with *Itgb1* shRNA. (H) GSEA analysis of TELSC-TO and TELSC-TO transfected with *Itgb1* shRNA based on “Establishment of cell polarity” and “Integrin pathway” geneset.

## EXPERIMENTAL MODEL AND SUBJECT DETAILS

### KEY RESOURCES TABLE

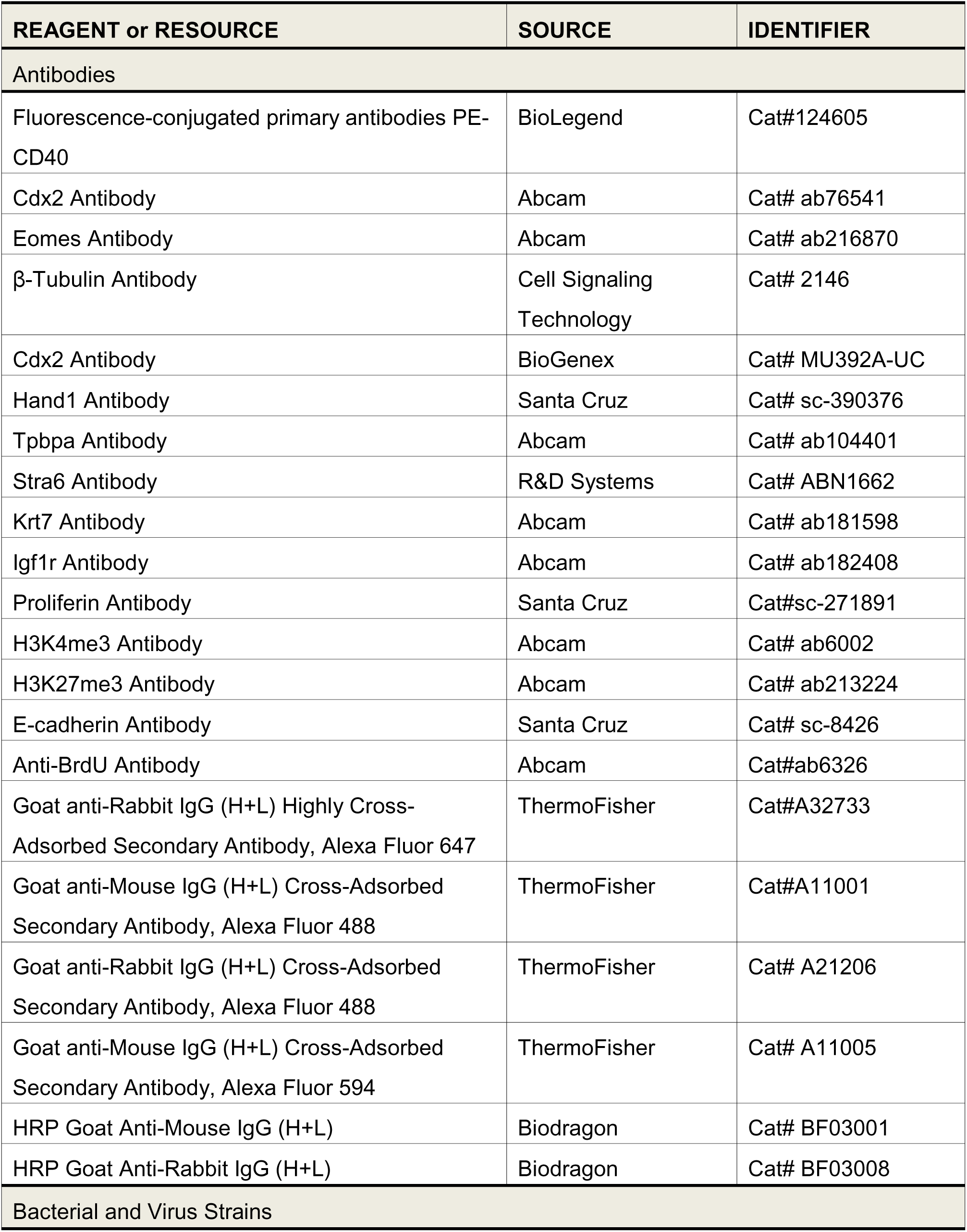
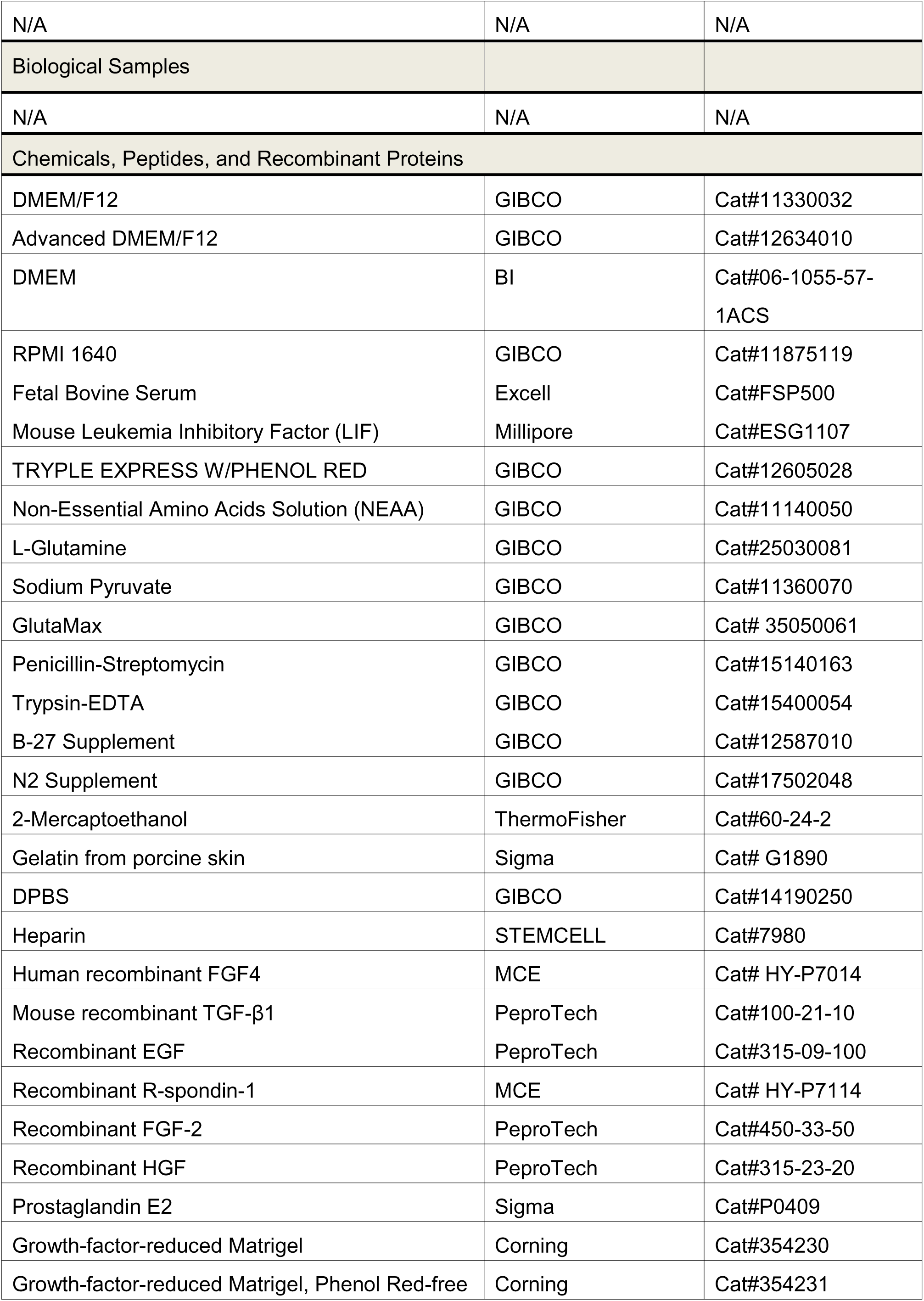
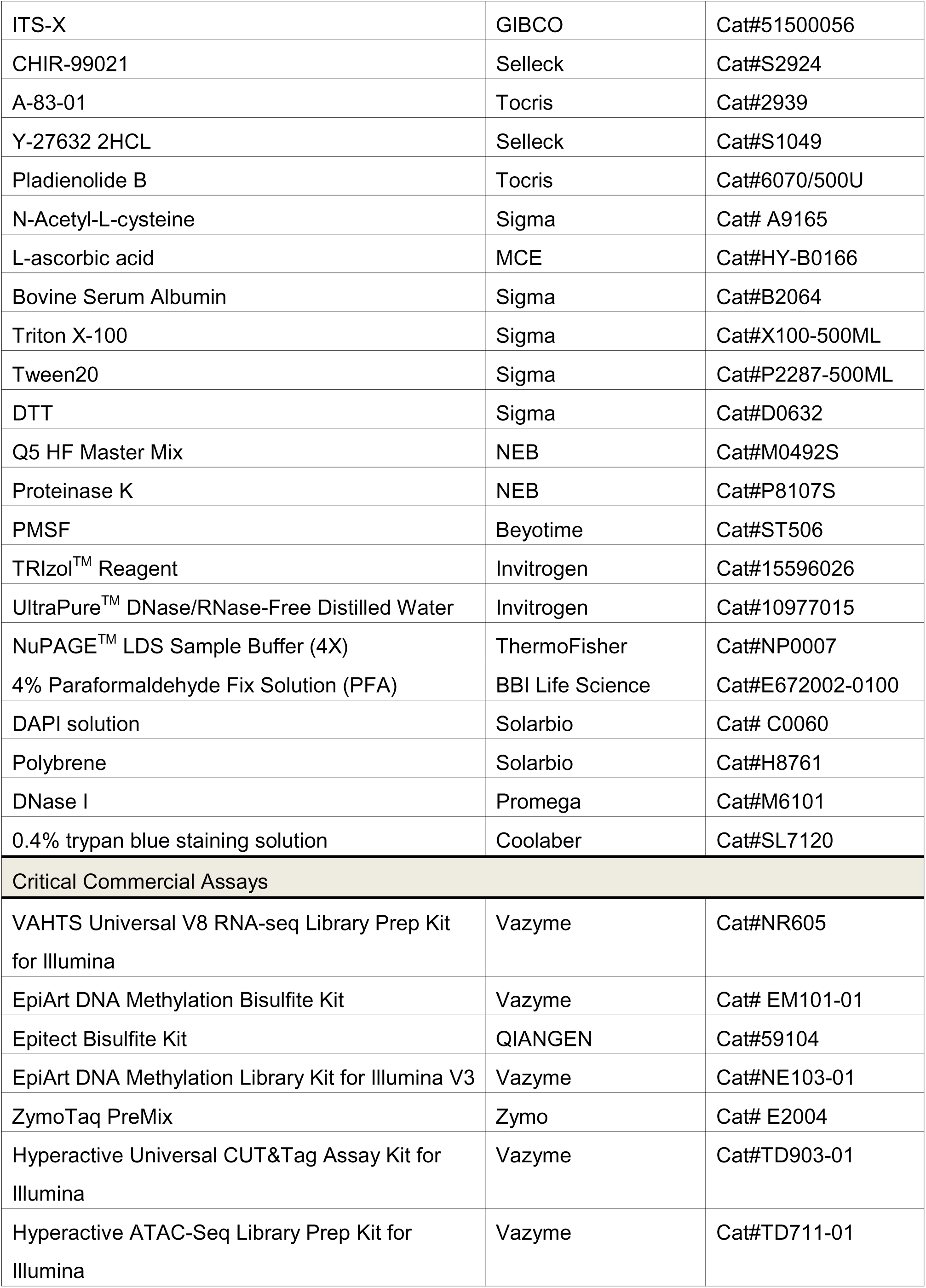
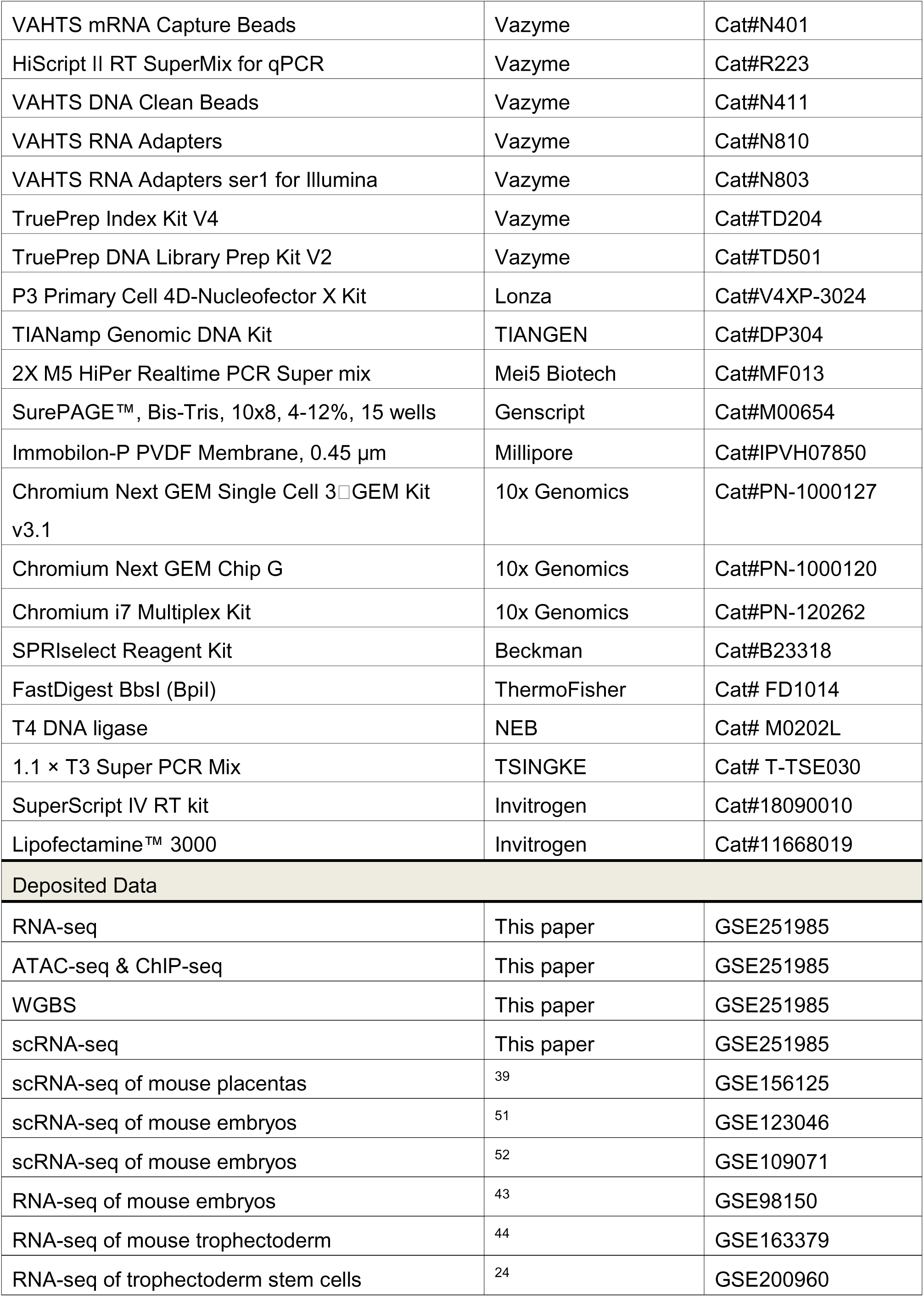
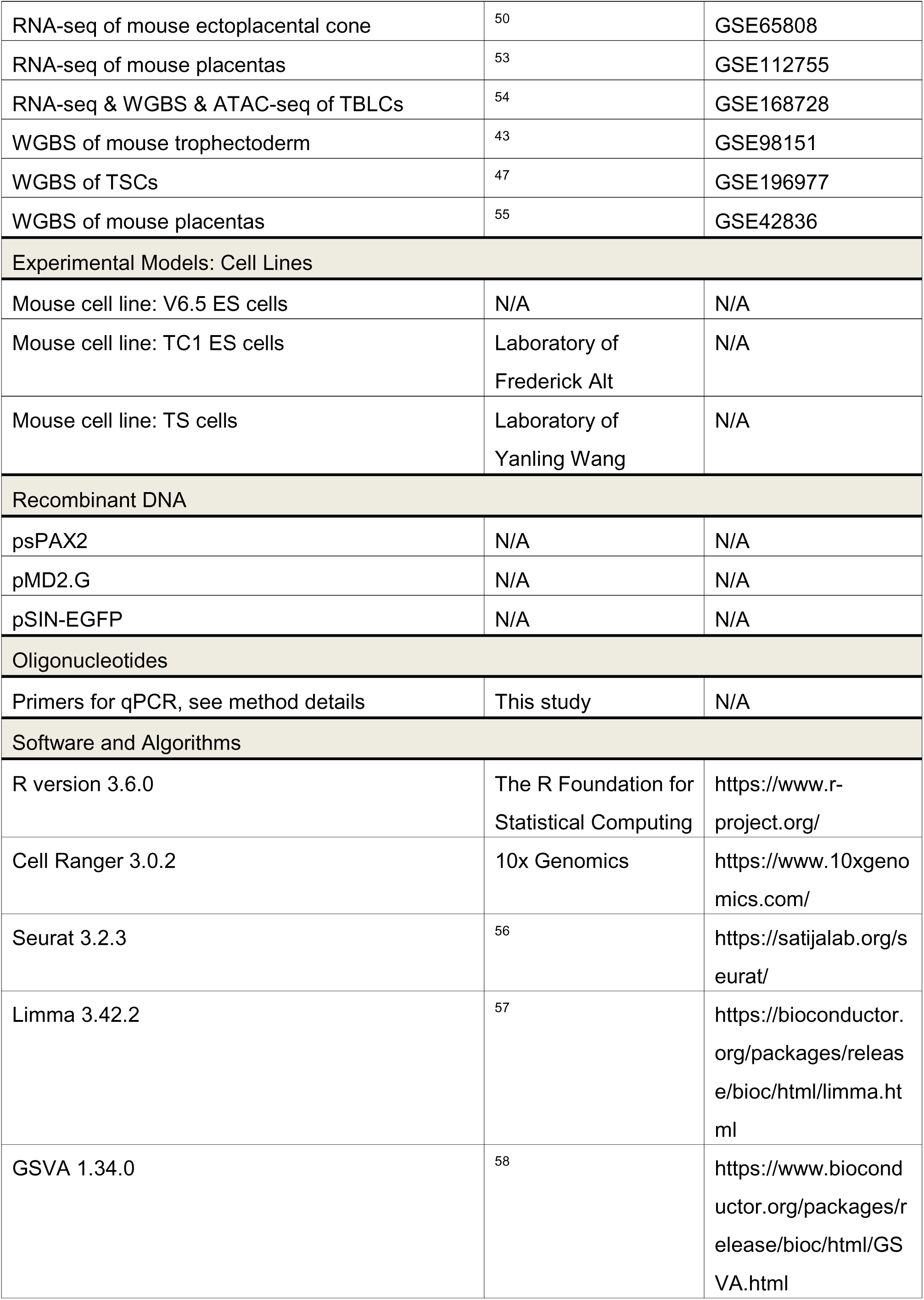
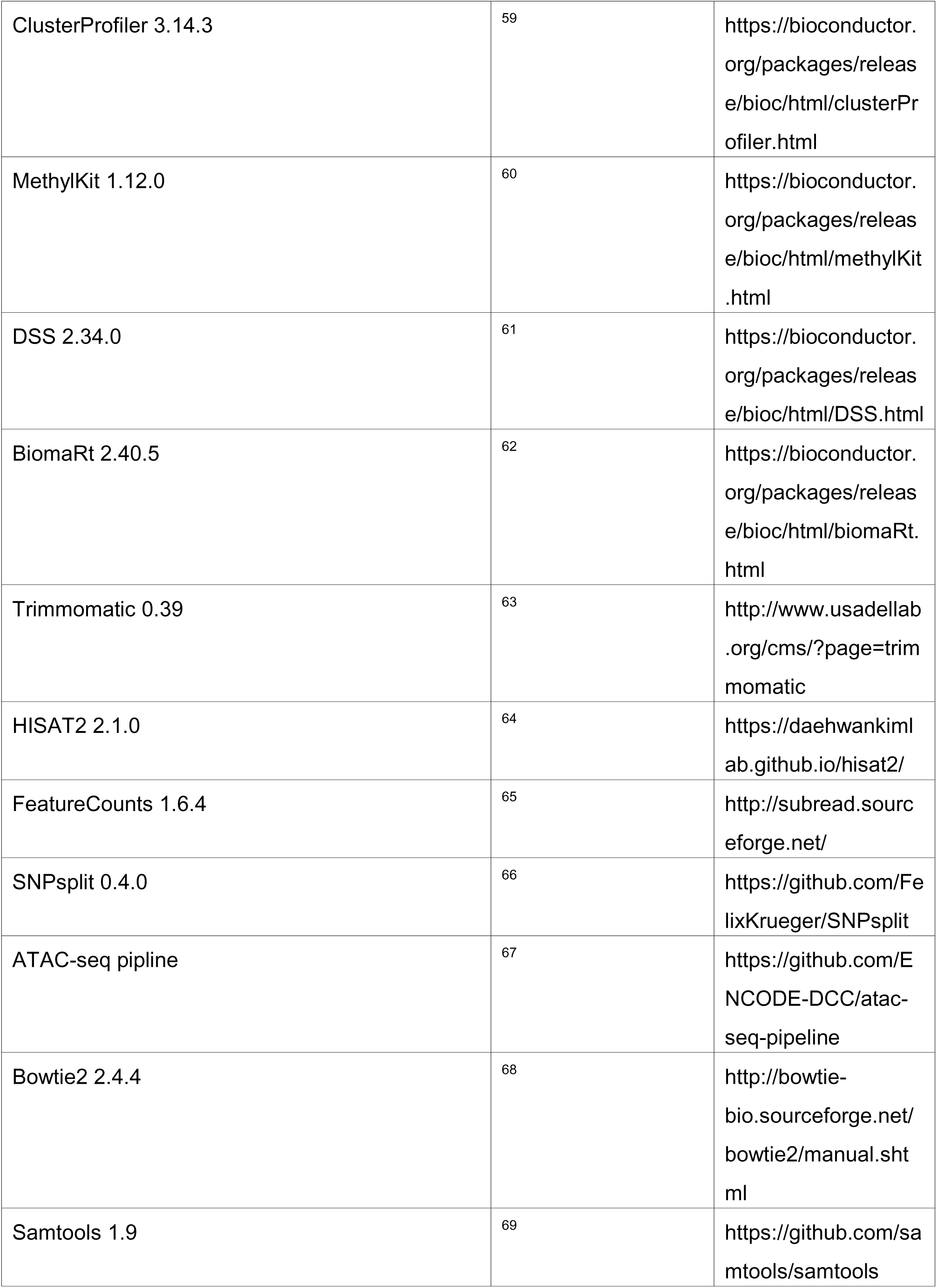
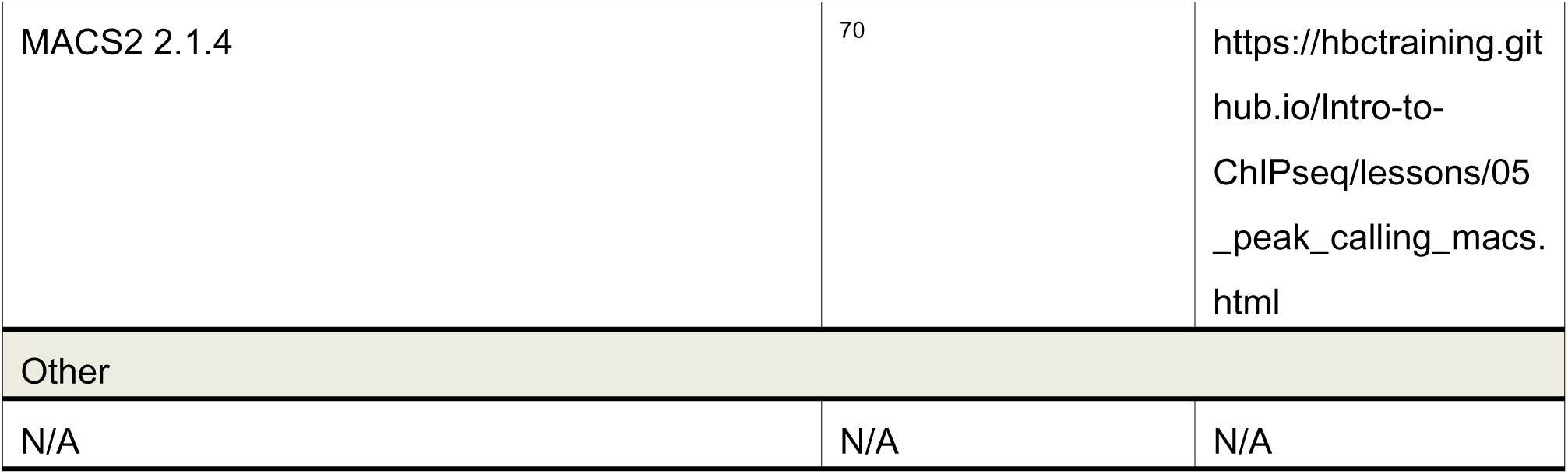

### RESOURCE AVAILABILITY

#### Lead contact

Further queries and reagent requests may be directed and will be fulfilled by the lead contact, Peng Du.

#### Data and code availability

The RNA-seq, scRNAseq, WGBS, ATAC-seq and ChIP-seq data generated in this study are available at GEO: GSE251985. All samples generated during this study have been summarized in **Table S7**.

## EXPERIMENTAL MODEL AND SUBJECT DETAILS

### METHOD DETAILS

#### Cell culture

All cell lines were cultured under 20% O_2_ and 5% CO_2_ at 37°C, and the medium was refreshed daily. Cells were passaged every 2-3 days with 0.25% trypsin-EDTA (GIBCO, 15400054). All ESCs (V6.5, TC1, M3, female ESCs and reporter cell lines) were cultured on mitomycin-treated mouse embryonic fibroblast (MEF) feeder layers or 0.1% gelatin (Sigma, G1890), in serum/LIF medium composed of DMEM (GIBCO, 11965092) supplemented with 20% fetal bovine serum (FBS) (ExCell, FCS500), 1% L-glutamine (GIBCO, 25030081), 1% penicillin-streptomycin (GIBCO, 15140163), 1% non-essential amino acids (NEAA) (GIBCO, 11140050), 1% sodium pyruvate (GIBCO, 11360070), 50 mM 2-mercaptoethanol (ThermoFisher, 60-24-2) and 1000 U/mL mouse leukemia inhibitory factor (mLIF) (Millipore, ESG1107). Feeder cells were seeded (5×10^4^ cells per cm^2^) onto 0.1% gelatin-coated dishes at least 12 hours before seeding ESCs. Feeder plates were used within 3 days.

All TSLs were cultured on Matrigel-coated plates, in 30% TS medium (RPMI 1640 (GIBCO, 11875119), 20% FBS, 1% GlutaMax (GIBCO, 35050061), 1% penicillin-streptomycin (GIBCO, 15140163), 1% sodium pyruvate (GIBCO, 11360070)) and 70% MEF-conditioned TS medium supplemented with 25 ng/ml human recombinant FGF4 (MCE, HY-P7014) and 1 μg/ml heparin (STEMCELL, 7980). For longer culture, the medium was changed to serum-free TX medium (Garreta et al., 2021; Kim et al., 2020; Rossi et al., 2018): F12/DMEM (GIBCO, 11330057), 64 mg/L L-ascorbic acid-2-phosphate magnesium, 1% penicillin-streptomycin, 1% sodium pyruvate, and 2% Insulin-Transferrin-Selenium-X (GIBCO, 51500056), supplemented with 25 ng/mL human recombinant FGF4, 1 μg/mL heparin, and 2 ng/mL mouse recombinant TGF-β1 (PeproTech, 100-21-10). TSLs were passaged every three days by incubation in TrypLE (GIBCO, 12605028) for 3 minutes, and the enzyme was inactivated by the addition of TS medium.

To generate TBLCs, ESCs were cultured in SLP medium (serum/LIF medium supplemented with 2.5 nM PlaB (Tocris, 6070) or 10 nM plaB (Kubaczka et al., 2014). For the first passage, cells were plated in serum/LIF medium, and the medium was changed to SLP medium the following day. For cryopreservation, cell pellets were resuspended in FBS/10% DMSO and stored at −80℃ or in liquid nitrogen.

#### Derivation of TELSC^em^ line from 8-cell embryos

Using a microscope, the zona pellucida of at least 40 eight-cell mouse embryos was digested with Tyrode’s solution at room temperature for approximately 5 minutes. Following digestion, embryos were transferred via mouth pipette to a gelatin-coated 48-well plate and cultured in TS medium. Initially, cell proliferation was slow, with visible cell aggregates appearing after approximately one week. These cells were then transitioned into TX medium for extended culture over at least five passages. The establishment of the TELSC^em^ line was confirmed through RNA sequencing and immunofluorescence staining.

#### Teratoma formation assay

MEFs, TSCs or TELSCs were digested into single cells, and approximately 5×10^6^ cells were collected and re-suspended in 100 μl of Matrigel. Next, the cells were subcutaneously injected into each side of 6- to 8-week-old immunodeficient NOG female mice. After growth for 1-2 weeks on average (day 10), all mice were euthanized via cervical dislocation and teratomas were extracted via surgical excision using scissors and forceps for further analysis.

For hematoxylin and eosin (H&E) staining and immunohistochemistry (IHC) staining, the teratomas were fixed overnight at 4°C in 4% PFA, embedded in paraffin and subsequently processed for staining and analysis. For scRNA-sequencing, the teratomas were cut into small pieces on ice and digested at 37°C with collagenase IV supplemented with 1 U/mL DNase for 30 min. Single cells were re-suspended in 0.04% BSA for the 10x Genomics Chromium platform.

#### Blastocyst chimera assay

All mouse embryo microinjection and transplantation experiments were performed at Beijing Vitalstar Biotechnology Laboratory. EGFP- or mCherry-labelled TELSCs and TSCs were digested into single cells using TrypLE, washed twice with cold PBS and resuspended in culture medium. The cell suspension were incubated on ice for at least 30 min before microinjection.

Before microinjection, mouse embryos at 8-cell or blastocyst stage were obtained from the oviducts of superovulated ICR mice with M2 medium. Collected embryos were then transferred to M2 droplets and cultured under 5% CO_2_ at 37°C covered with mineral oil until further processing.

To create chimeric placentas, 10–15 TELSCs or TSCs were microinjected into each 8-cell or blastocyst stage ICR mouse embryo, respectively. After microinjection, embryos were recovered in KSOM drops for 1–2 h. Then, 10–15 chimeric embryos were transferred to a surrogate mouse and harvested at the E10.5–E13.5 developmental stage for analysis.

Conceptuses were dissected on ice in PBS/10% FBS, and the fetus, yolk sac and placenta were separated from the conceptuses with fine pointed forceps and imaged using a stereo fluorescence microscope (Leica, M205 FCA) to localize EGFP^+^ cells. After washing with PBS, tissues were then minced into approximately 1 mm fragments on ice using fine-pointed forceps. Placentas were digested at 37°C for 30 min with collagenase IV (Gibco, 17104019) supplemented with 1 U/mL DNase (Promega, M6101). PBS with 10% FBS was added to stop the reaction, and cells were centrifuged at 800 rpm for 5 min at 4°C. After resuspension in PBS/0.3% BSA, and the percentage of EGFP+ cells in the digested tissues was analyzed by flow cytometry.

#### Flow cytometry

ESCs and TELSCs were digested into single cells using 0.25% trypsin-EDTA or TrypLE and resuspended in 2% FBS (diluted with PBS (BI, 02—023—1ACS)). The cells were washed once with 4 mL of 2% FBS, resuspended in 100 μL of 2% FBS, and incubated on ice for 10 minutes. Then, fluorescence-conjugated primary antibodies (PE-CD40, BioLegend, 124605) were added at predetermined optimum concentrations and incubated on ice for 20 minutes in the dark. The samples were washed 3 times with at least 2 mL of 2% FBS by centrifugation at 800 rpm for 3 minutes. The cell pellet was resuspended in 0.5 mL of 2% FBS, and DAPI solution (Solarbio, C0060) was added to exclude dead cells if necessary. The cell suspension was then filtered through 40 μm cell strainers. Fluorescence-activated cell sorting (FACS) or flow cytometric analysis was performed. Data analysis was performed using FlowJo software. FACS-enriched cells were centrifuged and plated on gelatin/feeder/Matrigel-coated dishes for further culture.

#### Western blotting

ESCs and TELSCs were lysed in cold lysis buffer (20 mM Tris-HCl (pH 7.6), 137 mM NaCl, 1 mM EDTA, 1% Triton X-100, 1.5 mM MgCl_2_) supplemented with 1 mM PMSF (Beyotime, ST506) and then mixed with 4 × NuPAGE™ LDS sample buffer (Invitrogen, NP0007) before incubation at 98°C for 10 minutes. Then, the protein samples were loaded for SDS-PAGE and transferred to PVDF membranes (Millipore, IPVH07850) using a Bio-Rad transfer apparatus. The membrane was blocked with 5% milk (diluted in TBS buffer containing 0.1% Tween-20 (TBST) at room temperature for 1 hour, followed by incubation with primary antibody overnight at 4°C. After 3 washes with TBST, the membranes were incubated with appropriate HRP-conjugated secondary antibodies at room temperature for 1 hour with shaking. After three washes in TBST, the membrane was imaged with an Amersham Imager 600 (GE Healthcare).The following primary antibodies were used: anti-CDX2 (Abcam, ab76541), anti-EOMES (Abcam, ab216870), anti-OCT4 (Abcam, ab18976), and anti-β-Tubulin (Cell Signaling Technology, 2146).

#### Derivation of trophoblast organoids from TELSCs

After FACS, TELSCs were centrifuged and re-suspended in an appropriate volume of growth factor-reduced Matrigel (Corning, 354230) on ice to generate 30 μL Matrigel containing 0.5-1.0×10^4^ cells. Drops (30 µL) were plated into each well of a 24-well culture plate, maintained at 37°C for 15 minutes and overlaid with 800 µL trophoblast organoid medium (TOM). Cultures were maintained in 5% CO_2_ in a humidified incubator at 37°C. The medium was replaced every 2 days. Small organoid clusters became visible by approximately day 7 and were collected for downstream experiments.

TOM is composed of Advanced DMEM/F12 (GIBCO, 12634-010), 100 × N2 supplement (Thermo, 17502048), 50 × B27 supplement minus vitamin A (Thermo, 12587010), 1.25 mM N-Acetyl-L-cysteine (SIGMA, A9165), 2 mM GlutaMax, 50 ng/mL recombinant EGF (PeproTech, 315-09-100), 1.5 µM CHIR99021 (Selleck, S2924), 80 ng/mL recombinant R-spondin-1 (MCE, HY-P7114), 100 ng/mL recombinant FGF-2 (PeproTech, 450-33-50), 50 ng/mL recombinant HGF (PeproTech, 315-23-20), 500 nM A83-01 (Tocris, 2939), 2.5 µM prostaglandin E2 (SIGMA, P0409), and 2 µM Y-27632 (Selleck, S1049). The medium was stored at 4°C for up to 1 week.

#### qRT-PCR

TRIzol reagent (Invitrogen, 15596026) was used to isolate total RNA. Briefly, 1 mL TRIzol reagent was added to the collected cells, and after 5 minutes of incubation at room temperature, 0.2 mL chloroform was added for phase separation and incubated for 2 minutes. Then the samples were centrifuged for 20 minutes at 14,000 × rpm at 4°C. The upper aqueous phase was transferred to a new 1.5 mL tube and 0.5 mL isopropanol was added to precipitate the RNA. After mixing, the samples were incubated for 1 hour at −20°C, and then centrifuged for 30 minutes at 14,000 × rpm at 4°C. The supernatant was discarded and the RNA pellets were washed with 75% ethanol. Finally, the RNA pellets were air-dried and dissolved in RNase-free water. For mRNA qRT-PCR analysis, 500 μg RNA was converted into cDNA via reverse transcription using HiScript IIQRT SuperMix for qPCR (Vazyme, R223). Then, 2× Real-time PCR Mix (Mei5 Biotech, MF013) and an Applied Biosystems SteponePlus Real-Time PCR System (Thermo Fisher) were used to quantify the cDNA in duplicate or triplicate. Gene expression was normalized to Gapdh. The primers used are listed in **Table S8**.

#### Immunofluorescence of cultured cell, organoids, and embryos

ESCs and TELSCs were grown on gelatin/Matrigel-coated glass coverslips. The cells were fixed with fresh 4% paraformaldehyde (PFA)/PBS for 20 minutes at room temperature, washed three times with PBS, and permeabilized in 0.2% Triton X-100/PBS for 10 minutes at room temperature. The cells were blocked with 3% BSA/PBS and incubated with the primary antibody diluted in 3% BSA/PBS overnight at 4℃. Cells were then washed three times with PBS, incubated with secondary antibodies for one hour at room temperature, washed in PBS for three times, mounted with DAPI and imaged with a confocal microscope (Dragonfly High Speed Spinning Disk Confocal Microscope).

Organoids were grown in 35 mm confocal dishes (Cellvis, D35-20-1-N), fixed in 4% PFA for 30 minutes at room temperature, washed three times in PBS, permeabilized for 30 minutes in 0.5% Triton X-100/PBS, washed in PBS for three times and blocked in 5% BSA/PBS for 40 minutes at room temperature. Primary antibodies were incubated in 5% BSA/PBS with 0.05% Triton at 4℃ overnight. Organoids were washed three times for 15 minutes in PBS and incubated for 3 hours at room temperature in 5% BSA/PBS with secondary antibodies. Organoids were mounted with DAPI and imaged with a confocal microscope (Dragonfly High Speed Spinning Disk Confocal Microscope).

The following primary antibodies were used: anti-CDX2 (Abcam, ab76541)/(BioGenex, MU392A-UC), anti-EOMES (Abcam, ab216870), anti-ELF5 (Santa Cruz, sc-376737), anti-GATA3 (Abcam, ab199428), anti-HAND1 (Santa Cruz, sc-390376), anti-TPBPA (Abcam, ab104401), anti-STRA6 (R&D Systems, ABN1662), anti-KRT7 (Abcam, ab181598), anti-IGF1R (Abcam, ab182408), anti-PROLIFERIN (Santa Cruz, 271891), anti-OCT4 (Abcam, ab181557), anti-SOX2 (Abcam, ab137385), anti-β-Tubulin (Cell Signaling Technology, 2146), and anti-KIP2 (Abcam, ab133531).

#### PolyA(+) RNA-seq

For sample preparation, 1 μg of total RNA and a VAHTS Universal V8 RNA-seq Library Prep Kit (Vazyme, NR605) were used according to the manufacturer’s instructions. Library samples were subjected to the Illumina HiSeq X Ten sequencing system. For the analysis of RNA-seq data, Trimmomatic software was used to trim the adapters. HISAT2 software was used for alignment to the mm10 reference genome with default parameters. The expression count matrix was obtained by FeatureCounts. Downstream analysis was performed with R (version 3.6.2).

#### Whole genome bisulfite sequencing (WGBS)

In total, 100 ng genomic DNA was used as input and was bisulfite-converted using the EpiArt DNA Methylation Bisulfite Kit (Vazyme, EM101-01). Bisulfite-converted DNA was used to construct the library with the EpiArtTM DNA Methylation Library Kit V3 (Vazyme, NE103-C2) according to the manufacturer’s instructions. Library samples were subjected to an Illumina Nova-seq 6000 sequencing system.

#### Bisulfite genomic sequencing (BSP)

1 μg of genomic DNA from ESCs, TSLs and TOs were processed for bisulfite sequencing analysis using the Epitect Bisulfite Kit (QIANGEN, 59104) according to the manufacturer’s instructions. PCR reactions were performed for all genes analyzed (including Elf5, Oct4, and Nanog) using ZymoTaq PreMix (Zymo, E2004).

#### CUT&Tag

A total of 5×10^4^ viable cells from ESCs, TSLs and TOs were used for CUT&Tag experiments with the Hyperactive Universal CUT&Tag Assay Kit for Illumina (Vazyme, TD903) according to the manufacturer’s instructions. Library samples were subjected to the Illumina HiSeq X Ten sequencing system.

#### ATAC-seq library preparation and sequencing

ATAC-seq was performed using the TruePrep DNA Library Prep Kit V2 for Illumina (Vazyme, TD501). A total of 5×10^4^ viable cells from ESCs, TELSCs and TOs were digested and collected by centrifugation at 800 rpm for 5 minutes. The cell pellets were resuspended in cold lysis buffer (1M Tris-HCl, pH 7.4, 5 M NaCl, 1 M MgCl_2_, 0.1% NP40, 0.1% Tween-20 and 0.01% digitonin) and incubated on ice for 3 minutes. Then, the cell pellets were washed with 1 mL of wash buffer (1 M Tris-HCl, pH 7.4, 5 M NaCl, 1 M MgCl_2_ and 0.1% NP40) by inverting the tube three times. The cells were collected by centrifugation at 500 rcf for 10 minutes, and the cell pellets were resuspended in 50 mL transposition mix according to the manufacturer’s instructions. Next, the samples were subjected to an Illumina Nova-seq 6000 sequencing system.

#### Single-cell RNA sequencing

Mouse TOs were dissociated at day 9 as described previously (Karvas et al., 2022). In brief, organoids were removed from Matrigel in cell recovery solution at 4°C for 30 minutes. Organoids were collected and washed in ice-cold HBSS. Activated Papain solution was added and incubated at 37°C for 10 minutes. 10% FBS in DMEM was added to finish the digestion. 10 ml of DNase I (Promega, M6101) was added to each tube and incubated at 37°C for 5 minutes. Samples were centrifuged at 4000 RPM for 30 seconds. The cells were resuspended in PBS+0.04% BSA. Single-cell libraries were constructed using the 10× Single-cell 3L Library & Gel Bead Kit v3.1 according to the manufacturer’s protocol.

### QUANTIFICATION AND STATISTICAL ANALYSIS

#### PolyA(+) RNA-seq data processing

All raw reads were first trimmed by Trimmomatic (version 0.39) ^63^ software to remove adapters and low-quality reads. Then, clean data were mapped to human genome (hg38) using HISAT2 (version 2.1.0) with default parameters. The count matrix of gene expression in each sample was generated by FeatureCounts (version 1.6.4) ^65^. After the genes were normalized to all mapped reads of each sample (CPM: count per million), gene expression changes were calculated based on the following formula: FC (fold change) = (CPMa+5)/(CPMb+5). We filtered significantly changed genes with different cutoffs, as indicated in the figure legends. Pheatmap R package was ultimately used to visualize the dynamic gene expression of samples. The published sequence fastq files were downloaded from different databases. RNA-seq of mouse embryos data (GSE98150), RNA-seq of mouse trophectoderm data (GSE163379), RNA-seq of mouse ectoplacental cone data (GSE65808), RNA-seq of mouse placentas data (GSE112755) and RNA-seq of TBLCs data (GSE168728) were obtained from GEO database. These data were also analyzed using the same methods described above.

#### Single-cell RNA-seq data processing

Single-cell transcriptome data were first applied to Cell Ranger (version 3.6.0) against hg38 genome reference to generate count matrix, which of the column was cell name and the row was gene. The downstream analysis was performed in Seurat (version 3.1.0) R package ^56^. Then, we merged the single-cell data with published data (Jiang et al., 2023) directly. The cells from different samples can be detected in each lineage and mixed very well in each cluster indicating that the batch effect did not affect the cell clustering. The read count data were normalized by NormalizeData with “LogNormalize” method. We chose 1-30 PCs after RunPCA function to perform RunUMAP function and cell clusters were investigated by a shared nearest neighbor (SNN) modularity optimization with FindClusters function with parameter “resolution = 0.4”. Next, we used function “FindAllMarkers” to identify DEGs. Genes with “min.pct = 0.5, logfc.threshold = 0.25” were considered significantly different.

#### ATAC-seq data processing

ATAC-seq data were firstly trimmed by Trimmomatic software (version 0.39). Bowtie2 (version 2.4.4) ^68^ was used to align the clean reads to mm10 reference genome “-X 1000 –mm -local”. Picard was used to remove duplicates and we only remained the high-quality reads. Then, after calling the peaks by MACS2, we used the merge and multicov commands in bedtools to extract the count of the peaks in all the samples and bdgdiff function in MACS2 was used to normalize the count matrix and calculate the differential peaks with parameters “-g 100 -l 50 -C 2”. ATAC-seq data of mTBLC were downloaded from GEO database (GSE168728), and analyzed using the same method.

#### ChIP-seq data processing

ChIP-seq data were firstly trimmed by Trimmomatic software. Bowtie2 (version 2.4.4) ^68^ was used to align the clean reads to mm10 reference genome with parameters “--end-to-end --very-sensitive --no-mixed --no-discordant --phred33 -I 10 -X 700”. We used MACS2 (version 2.1.4) ^70^ to call peaks and differential peaks were detected by macs2 bdgdiff command.

#### Principal component and unsupervised hierarchical clustering analysis

We used PCA function in FactoMineR (version 1.42) and hclust function to perform PCA and clustering analysis, respectively. Due to the batch effects in different projects, Combat function in sva (version 3.34.0) ^71^ R package was used to remove the batch effect between RNA-seq data obtained from different studies.

#### Gene set variation analysis (GSVA)

Gene set variation analysis was carried out using the GSVA (version 1.32.0) R package.

#### Gene set enrichment analysis (GSEA)

GSEA was performed using row counts with GSEA software (version 4.1.0). The enrichment score was calculated by GSEA function in cluster Profiler R package.

#### Gene ontology annotation

ClusterProfiler (version 3.14.3) R package was used for Gene Ontology (GO) term analysis. GO terms with p-value < 0.05 were defined as significant process.

#### Pseudotime trajectory analysis

Pseudotime trajectories were constructed using Monocle2 (2.22.0). Only genes with an average expression greater than 3 were retained for the analysis. The “FindVariableFeatures” function was used to identify highly variable genes for ordering cells. The “plot_cell_trajectory” function was used to visualize results annotated with cell type information.

#### Alignment track visualization

The visualization of different tracks was performed using IGV.

## Supplemental Information

Table S1. Lists of differentially expressed genes in TBLCs, TELCs, TELSCs, TESC (Seong et al., 2022) and TSC, related to Figure 1

Table S2. Lists of differentially expressed genes in TE3.5-ExE6.5 cells used in GSVA, related to Figure 1K

Table S3. Lists of differentially expressed genes in MEF-, TSC- and TELSC derived teratoma, related to Figure S4E

Table S4. Lists of differentially expressed genes in in MEF-, TSC- and TELSC derived teratoma used in GSVA, related to Figure 4F

Table S5. Marker genes for each defined trophoblast cell type (ExE-like, LaTP, SpT, SynTI, SyntII, GlyT, S-TGC, P-TGC), related to Figure 6

Table S6. Lists of differentially expressed genes in TELSC, TELSC-derived organoids at different time points and organoids reported by Mao et al.,, related to Figure 7C

Table S7. Summary of all sequencing data in this study

Table S8. Lists of primers and sgRNA sequences used in this study

